# Stochastic semi-supervised learning to prioritise genes from high-throughput genomic screens

**DOI:** 10.1101/655449

**Authors:** Dimitrios Vitsios, Slavé Petrovski

**Affiliations:** Centre for Genomics Research, Discovery Sciences, BioPharmaceuticals R&D, AstraZeneca, Cambridge, UK

## Abstract

Access to large-scale genomics datasets has increased the utility of hypothesis-free genome-wide analyses that result in candidate lists of genes. Often these analyses highlight several gene signals that might contribute to pathogenesis but are insufficiently powered to reach experiment-wide significance. This often triggers a process of laborious evaluation of highly-ranked genes through manual inspection of various public knowledge resources to triage those considered sufficiently interesting for deeper investigation. Here, we introduce a novel multi-dimensional, multi-step machine learning framework to objectively and more holistically assess biological relevance of genes to disease studies, by relying on a plethora of gene-associated annotations. We developed *mantis-ml* to serve as an automated machine learning (AutoML) framework, following a stochastic semi-supervised learning approach to rank known and novel disease-associated genes through iterative training and prediction sessions of random balanced datasets across the protein-coding exome (n=18,626 genes). We applied this framework on a range of disease-specific areas and as a generic disease likelihood estimator, achieving an average Area Under Curve (AUC) prediction performance of 0.85. Critically, to demonstrate applied utility on exome-wide association studies, we overlapped *mantis-ml* disease-specific predictions with data from published cohort-level association studies. We retrieved statistically significant enrichment of high *mantis-ml* predictions among the top-ranked genes from hypothesis-free cohort-level statistics (p<0.05), suggesting the capture of true prioritisation signals. We believe that *mantis-ml* is a novel easy-to-use tool to support objectively triaging gene discovery and overall enhancing our understanding of complex genotype-phenotype associations.

## Introduction

The explosion of genomic data in the last decade has offered the scientific community a unique opportunity to elucidate the functionality of the building blocks of the human genome. A major area of focus has been decoding the clinical relevance of protein-coding genes, which are directly associated with cell stability, development but also cell proliferation, pathogenicity and disease. Given the vast interrogation and elucidation of the protein-coding genome, the global research community has generated an extended amount of resources with regards to tissue-specific gene expression, intolerance to genetic variation, model organism function and various other diverse types of annotation. Additionally, it is now evident that complex phenotypes, such as disease, are not represented by the variability of a single data type (e.g. expression in tissue or GWAS results) but rather require the combination of a multitude of data types and resources that describe multiple aspects of the phenotype at different dimensions^1,2,3^.

At the same time, tens-to-thousands of genes have been identified as essential contributors to complex diseases. The underlying biology for these diseases is complex and current knowledge provides a limited view of the full collection of disease-associated genes. We sought to explore this issue by leveraging the rich collection of gene-associated annotations to identify patterns shared among genes associated with a disease and leverage those patterns to predict putative genes of interest that may also be disease-associated.

We applied this framework to a number of disease areas, each time harvesting diverse types of information per gene, including gene expression^4^, human disease literature^5^, mouse phenotypes^6^, proteomic^7^, interactome^8^ and genic metrics of human-lineage purifying selection^9,10,11^. Our machine learning framework is implemented in a disease agnostic manner so that it can be applied to any disease area, given a sufficient starting set of known disease-associated genes and is demonstrably relevant when applied as a tool to orthogonally triage the results of exome-wide association statistics. To illustrate this, we have applied *mantis-ml* to three broad disease groups where exome-wide association statistics have been published: Amyotrophic Lateral Sclerosis (ALS), Chronic Kidney Disease (CKD) and Epilepsy.

## Results

### Automated feature pre-processing

*mantis-ml* has been developed as an automated machine learning (AutoML) framework to enable learning from an arbitrary set of gene-associated features (**Suppl. Fig. 1**). We collated data from a diverse set of gene-annotation sources (**Fig. 1a** and **Suppl. File 1**), classified into three categories: generic resources (disease/tissue agnostic), resources filtered by tissue and finally disease-specific features. Given a set of user-specified query terms relating to a tissue and/or disease of interest the data compilation and cleaning is performed automatically. This includes filtering of highly correlated features (default threshold: Pearson’s r ≥ 0.8) and features with high ratio of missing or undefined data (default missing data threshold=0.25). Missing/undefined data are imputed across all features, either with zero or the median value depending on the feature type (see Methods).

**Fig. 1.**
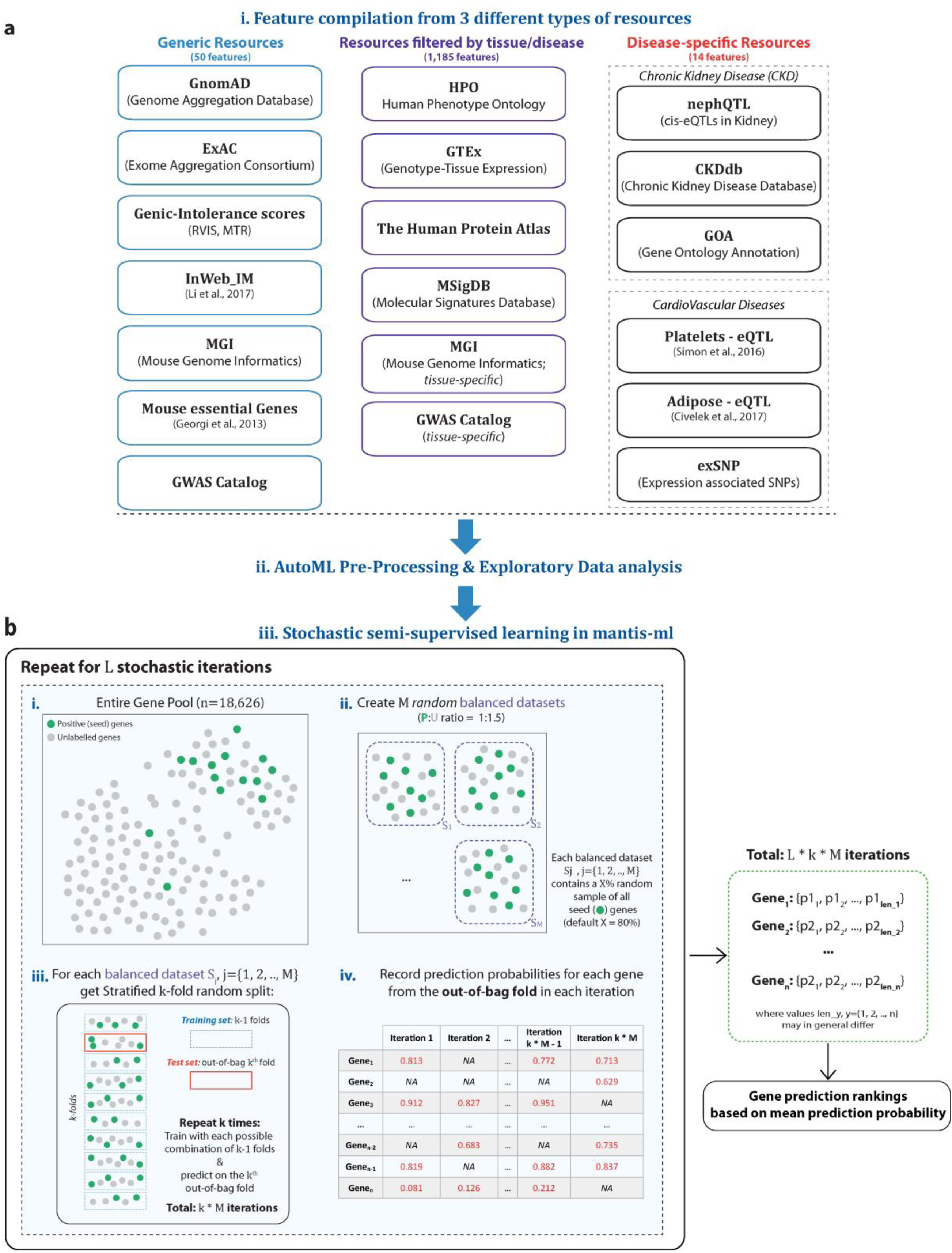
Generic overview of *mantis-ml* workflow. **a.** Data resources used by *mantis-ml* for feature extraction. Three data type resources are integrated: generic (i.e. non-tissue/disease specific), filtered by tissue-specific and also filtered by disease-specific features (currently including disease-specific features for Chronic Kidney Disease and Cardiovascular Disease). All features are compiled automatically based on user-provided disease-associated query terms and pre-processed, ready to be fed for the learning tasks. **b.** Illustration of the stochastic semi-supervised approach followed by *mantis-ml* over *L* iterations: i) positive (seed) genes are annotated using the Human Phenotype Ontology (static for each stochastic iteration), ii) the entire gene pool is split into random balanced sets, each of them including a random sample (default: 80%) of seed genes, iii) each balanced dataset is split with stratified *k*-fold, training is performed for each combination of *k-1* folds followed by prediction on the *k^th^* out-of-bag fold each time, iv) prediction probabilities are aggregated for each gene across all *L * k * M* iterations.

Furthermore, *mantis-ml* generates an automated set of visualisations for exploratory analysis, including heatmaps for pairwise feature correlations (prior and post feature filtering), missing data ratios across features and the distribution of numerical and/or categorical variables across the known and unlabelled genes from the entire gene pool (**Suppl. Fig. 2, 3**). Additionally, *mantis-ml* automatically performs dimensionality reduction on the original feature set using Principal Component Analysis (PCA), t-distributed Stochastic Neighbouring Embedding^12^ (t-SNE) and Uniform Manifold Approximation and Projection^13^ (UMAP) to allow for visualisation of the original high-dimensional space into two dimensions (**Suppl. Fig 4**). These visualisations aim to highlight any evident and/or trivial segregation of the known disease-associated genes from the unlabelled ones based on either pairwise relationships between the features or any linear and non-linear relationships captured by the first two or three projected vectors of the higher-dimensional feature space into the PCA, t-SNE and UMAP transformation spaces. However, the complexity underlying each of the diseases under study imposes the exploration of high-dimensional interactions between all features to elucidate the complex mechanisms that primarily drive pathogenicity in each case.

### Stochastic semi-supervised learning for gene prioritisation

*mantis-ml* seeks to uncover any feature patterns among a collection of known positive-labelled disease-associated genes to then prioritise novel genes that share a highly similar feature profile with the known disease genes. This problem falls into the broader machine learning area of positive-unlabelled learning, a semi-supervised learning technique where the only labelled data points available are positive. This is important in this context as we often have insufficient information about which genes among the remainder of the genome are definitively not associated with that disease (i.e., true negatives). There are several approaches aiming to solve positive-unlabelled problems, the most popular being: a) to treat unlabelled data as negative and perform learning with a standard classifier^14^, b) the use of bootstrap and bagging to iteratively train on random samples of positive and unlabelled data and predict on out-of-bag unlabelled data^15^ and c) use two-step approaches in which the first step tries to identify a confident set of negative points among the unlabelled set and then continues learning with a standard classifier^16^.

The positively-labelled genes for the gene prioritisation are retrieved by *mantis-ml* from the Human Phenotype Ontology^5^ (HPO) based on user-provided inclusion and exclusion query terms that are relevant to a particular disease (see Methods). The HPO component identifies seed genes based on the documentation of gene-disease association in OMIM and additional clinical terminology curated by clinicians participating in regular workshops hosted by the HPO team. An important peculiarity for this problem derives from the fact that in most diseases, the overall set of known data points (positive labels) is typically in the range of approximately 10^2^-10^3^ genes. This makes the entire gene set (n=18,626) highly imbalanced, in terms of the overall population of positive and unlabelled data points. Additionally, the entire gene space is finite and practically already known so we are not bound by the usual machine learning requirement to train a model that can generalise well on unseen data. Our end goal is to rank the entire gene set with respect to a diverse pool of disease groups, implying that our predictions should not be biased by a single training dataset of a subset of genes assumed to be representing the global distribution of all genes. As a result, and in combination with the lack of a well-defined negative set, training a model sufficiently generalisable to then predict on a test set would not be ideal in this context.

To address this issue, we constructed a novel gene prioritisation framework that is based on a variation of two of the positive-unlabelled approaches suggested above (a, b): a stochastic semi-supervised learning technique (*L* iterations) across multiple random balanced datasets from the entire gene set (**Fig. 1b** and **Suppl. Fig 1b**) with iterative predictions on out-of-bag data. In each stochastic iteration, we create a random partitioning of the unlabelled gene space to form *M* balanced datasets, with positive-to-unlabelled points ratio equal to 1:1.5. Each balanced dataset contains a random *X*% sample of the positive (seed) genes (default *X*=80%) to reduce bias induced by using the entire positive gene pool in each training task. The unlabelled data are treated as negative, since training on positive and unlabelled data in general gives scores proportional to the ones retrieved by training on positive and negative data^14^. We then perform stratified *k*-fold split on each balanced dataset (default *k*=10) and then train with a standard classifier for each possible combination of *k-1* folds (training set) followed by prediction each time on the out-of-bag *k^th^* fold (test set). This process is performed *k* times over each balanced dataset and upon each training cycle, prediction probabilities are retrieved only for the genes belonging to the respective test set (out-of-bag *k^th^* fold).

The entire procedure is repeated for *L* iterations, each one leading to a random set of balanced sets to allow inclusion of each gene into out-of-bag sets over multiple times and subsequently lead to more unbiased and robust results. *mantis-ml* does not define a static underlying model but prioritises all genes based on the probability prediction they have achieved over multiple iterations that have grouped them into random balanced gene groups. Eventually, we aggregate the prediction probabilities assigned to each gene member from out-of-bag sets (either positive or unlabelled) across all *L * k * M* iterations. This forms for each gene a probability distribution regarding association of a gene with the disease under examination. The final predictions are ranked based on the mean of their probability distribution (results equivalent with median, Pearson’s r > 0.9996, p<2.2×10^−308^).

### Classifier benchmarking for positive-unlabelled learning

We tested seven different classifiers to be used during positive-unlabelled learning for each balanced dataset: Random Forest, Extra Trees (Extremely Randomised Trees – a variation of Random Forest), Gradient Boosting, Extreme Gradient Boosting (XGBoost), Support Vector Classifier (SVC), Deep Neural Networks (DNN) and a Stacking (Ensemble) classifier with four base classifiers (Random Forest, Extra Trees, Gradient Boosting and SVC) followed by a DNN in the second layer (see Methods). We first fine-tuned each classifier separately using a single random balanced dataset from the CKD disease example with 10-fold Cross-Validation and performing Grid Search over a finite parameter space (see Methods). We then benchmarked all classifiers by assessing their AUC performance on the same set of ten random balanced datasets with 10-fold Cross-Validation. All classifiers performed comparably (average AUC: 0.831-0.850) with Random Forest and Extreme Gradient Boosting ranking as the top two classifiers, with mean AUC equal to 0.850 ± 0.021 and 0.848 ± 0.021, respectively (**Fig. 2** and **Suppl. Fig. 5**). Given the comparable performances across classifiers, we cannot pick yet a single classifier that outperforms the rest in the problem of gene prioritisation with positive-unlabelled learning. Thus, we are going to apply all classifiers to each disease example examined in this work and then select the best performing one within each disease example based on the average AUC scores achieved.

**Fig. 2.**
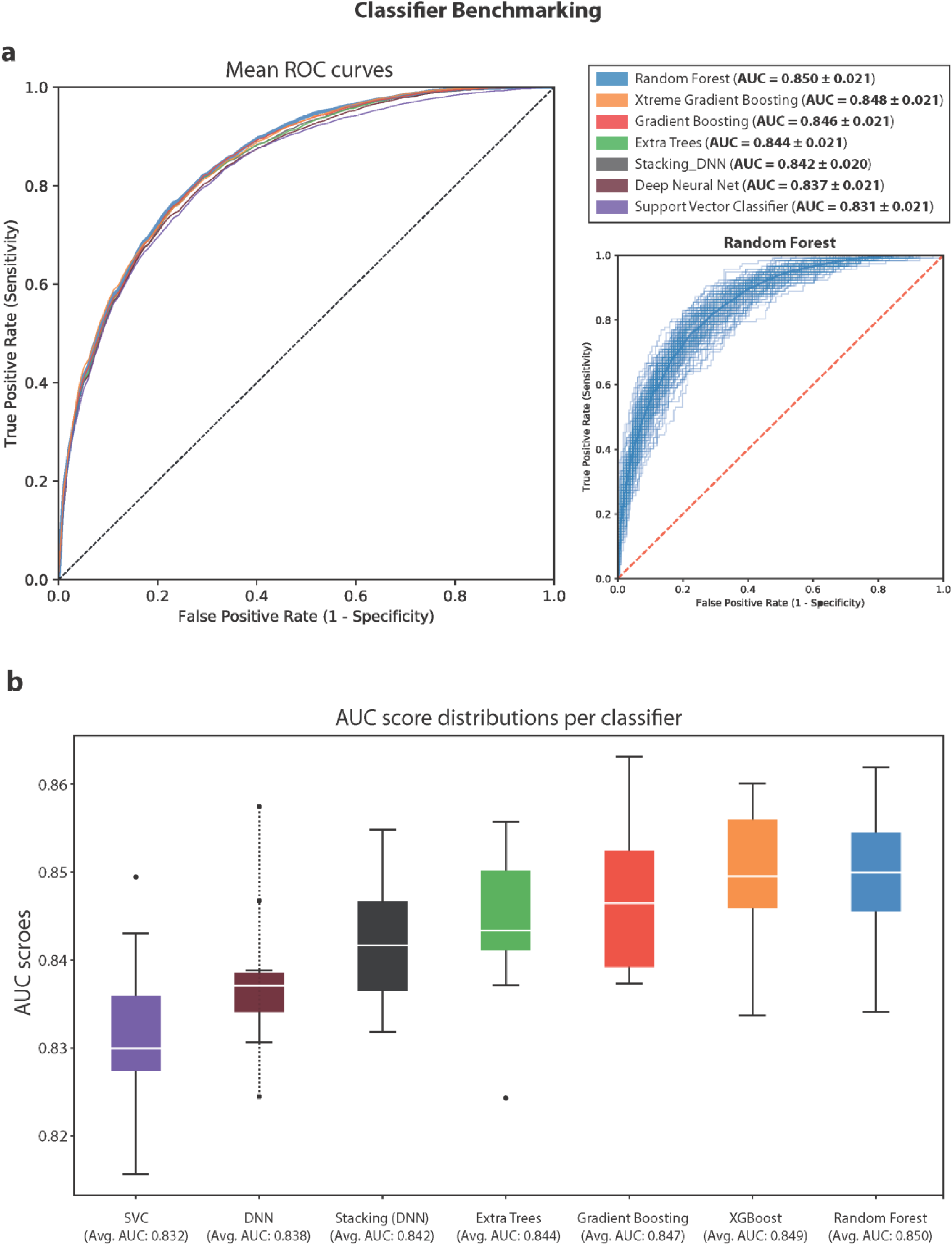
Benchmarking of seven different classifiers during the positive-unlabelled learning step of *mantis-ml*: Random Forest, Extra Trees, Gradient Boosting, Extreme Gradient Boosting (XGBoost), Support Vector Classifier (SVC), Deep Neural Network (DNN) and a Stacking (Ensemble) classifier with four base classifiers (Random Forest, Extra Trees, Gradient Boosting and SVC) followed by a DNN. *mantis-ml* was run on 10 random balanced datasets with 10-fold Cross-Validation based on the Chronic Kidney Disease example. a) Mean ROC curves from stochastic Positive-Unlabelled learning using one of the seven classifiers. ROC curves from all runs are also shown for the best performing classifier during benchmarking (Random Forest). b) Distribution of AUC scores across the seven classifiers tested. All classifiers showed comparable performance (AUC:0.83-0.85) with tree-based methods ranking at the top.

### Application on three disease examples: Amyotrophic Lateral Sclerosis, Chronic Kidney Disease, Epilepsy and

We have applied *mantis-ml* on three major disease categories with complex underlying disease mechanisms: Amyotrophic Lateral Sclerosis (ALS), Chronic Kidney Disease (CKD) and Epilepsy (Genetic Generalised Epilepsy). We selected these for demonstration of applied utility on hypothesis-free exome-wide association statistic results where we have previously published the full exome results. The positively labelled gene set for each disease was selected based on a user-defined curated dictionary of inclusion and exclusion terms used for automated querying and extraction from the Human Phenotype Ontology (HPO). Tissue specific and disease-relevant features were automatically extracted based on the same query terms applied to the *mantis-ml* integrated knowledgebase (see Methods). No selection of positive-labelled genes, tissue and disease-relevant features require human curation beyond provision of the user-defined curated dictionary of inclusion and exclusion terms. The total number of known (seed) positively-labelled genes found to be associated with each disease based on HPO where 587, 864 and 77 for CKD, genetic generalised epilepsy and ALS, respectively. We then applied all seven classifiers used during benchmarking across *L*=100 stochastic iterations for each disease-specific positive-unlabelled learning task. The total number of training/test tasks performed across an equivalent number of random balanced gene samples with cross-validation was 25,000, 17,000 and 79,500 for CKD, Epilepsy and ALS, respectively. These sizes are inversely proportional to the number of seed genes in each case, which directly affects the size of constructed balanced datasets across the entire gene pool.

All classifiers except for Stacking showed comparable performance when applied to the entire gene set for each disease case. The Stacking classifier consistently demonstrated slightly lower performance compared to the rest of classifiers (on average 0.05 lower AUC score). This is somewhat expected since one of Stacking’s most notable properties is to smooth out predictions from its base classifiers. This means that predictions supported by most of its base classifiers are more likely to survive in the end thus potentially lowering the total number of correctly identified genes, which is however likely to be more robust as compared to the predictions from each individual classifier due to its conservative nature.

XGBoost and Random Forest had the best performance in CKD (average AUC: 0.846) followed by Gradient Boosting and Extra Trees (average AUC: 0.843, 0.839, respectively), with some of the most well-established CKD genes (*PKD1*, *PKD2*, *COL4A1*, *COL4A3*, *COL4A4* and *COL4A5*) ranking in the top 0.2-0.7% of all genes (**Suppl. Fig. 6**). Notably, Extra Trees classifier ranks *PKD1* and *PKD2* in positions 5 and 27 among all ∼18K genes. An aggressive prediction probability threshold of 0.5 has been used to classify genes as either predicted known or novel. We observe high concordance between the predictions from all classifiers with regards to known disease genes. Specifically, 280 known genes (47.7% of all known genes) have been identified by all classifiers and another 64 genes (11% of all known genes) by at least five classifiers. In terms of the novel disease-associated genes, again the largest group of predicted genes-of-interest (n=1,300) has been predicted by all classifiers, while DNN yielded the highest number of novel predictions identified solely by this classifier. However, each classifier calculates their predictions with a ranking score (prediction probability) which immediately provides a prioritisation scheme for the extracted gene predictions. Having no further knowledge to validate *mantis-ml* predictions at this stage, we choose to consider the gene rankings of the classifier with the highest average and individual AUC scores (XGBoost) in that case as the default *mantis-ml* prioritisation scheme for CKD (**Suppl. File 2; Table S1**). We found that the *mantis-ml* gene prediction rankings were significantly correlated when comparing XGBoost and Random Forests (Pearson’s *r* = 0.976; p < 2.2 x 10^−308^, **Suppl. Fig. 20**), further demonstrating the robustness of the predictions beyond the choice of classifier.

Moving on to the *mantis-ml* performance on the genetic generalised epilepsy example, we observe again XGBoost, Random Forest, Gradient Boosting and Extra Trees performing best among all classifiers, ranked in descending order of average AUC scores: 0.821, 0.818, 0.816 and 0.808 (**Suppl. Fig. 7**). We can also observe the preponderance of 360 known genes (41.7% of all known genes) being predicted by all seven classifiers followed by another 67 genes (7.7% of all known genes) predicted by at least six classifiers (based on prediction probability threshold of 0.5). Additionally, around 1,600 novel genes have been suggested by all classifiers, while Stacking and DNN provide the highest numbers of novel gene predictions exclusively identified by each of them. We provide as the default gene ranking for Epilepsy the *mantis-ml* predictions acquired when using XGBoost as the standard classifier during positive-unlabelled learning, based on its best AUC performance among all classifiers (**Suppl. File 2; Table S2**).

Finally, with regards to *mantis-ml* predictions on ALS (**Suppl. Fig. 8**), Extra Trees followed by XGBoost, SVC and Random Forest showed the best performance (average AUC scores: 0.814, 0.805, 0.801 and 0.798). Moreover, 31 known (40.1% of all known genes) and 1,500 novel genes were predicted by all classifiers with Stacking and DNN again providing the highest number of novel predictions with support by a single classifier (based on prediction probability threshold of 0.5). We also observe in this disease example that a smaller number of seed genes (77) is accompanied by a slight drop in average AUC scores. This implies that *mantis-ml* performance is directly dependent on confidence from an increased presence of known genes, able to focus on broader patterns related with a disease that can then identify genes with similar feature profiles as known disease genes. We also provide the Extra Trees ranking scores as the default *mantis-ml* predictions for ALS (**Suppl. File 2; Table S3**).

We then sought to examine the importance of the seed genes set for the prediction performance of *mantis-ml*. We thus tested the prediction probabilities across all seed (positively-labelled) genes in the three disease examples (CKD, Epilepsy and ALS) when the seed gene set is selected based on either the HPO annotation or randomly (**Fig. 3a**). The size of seed gene sets varies in these disease examples (CKD: 587, Epilepsy: 864 and ALS: 77) allowing also exploration of the importance of seed gene lists of varying length. We observed that in all three diseases, the prediction probabilities for seed genes when using a real seed gene set are skewed towards a probability of ‘1.0’ while the respective distribution acquired when using random seed genes of same length is almost uniformly distributed across the entire probability spectrum (0.0-1.0). In all cases, the probability distributions significantly differ between real and random seed genes (Mann-Whitney U P-value=3.81×10^−87^, 2.34×10^−87^, 1.02×10^−08^ for CKD, Epilepsy and ALS, respectively).

**Fig. 3.**
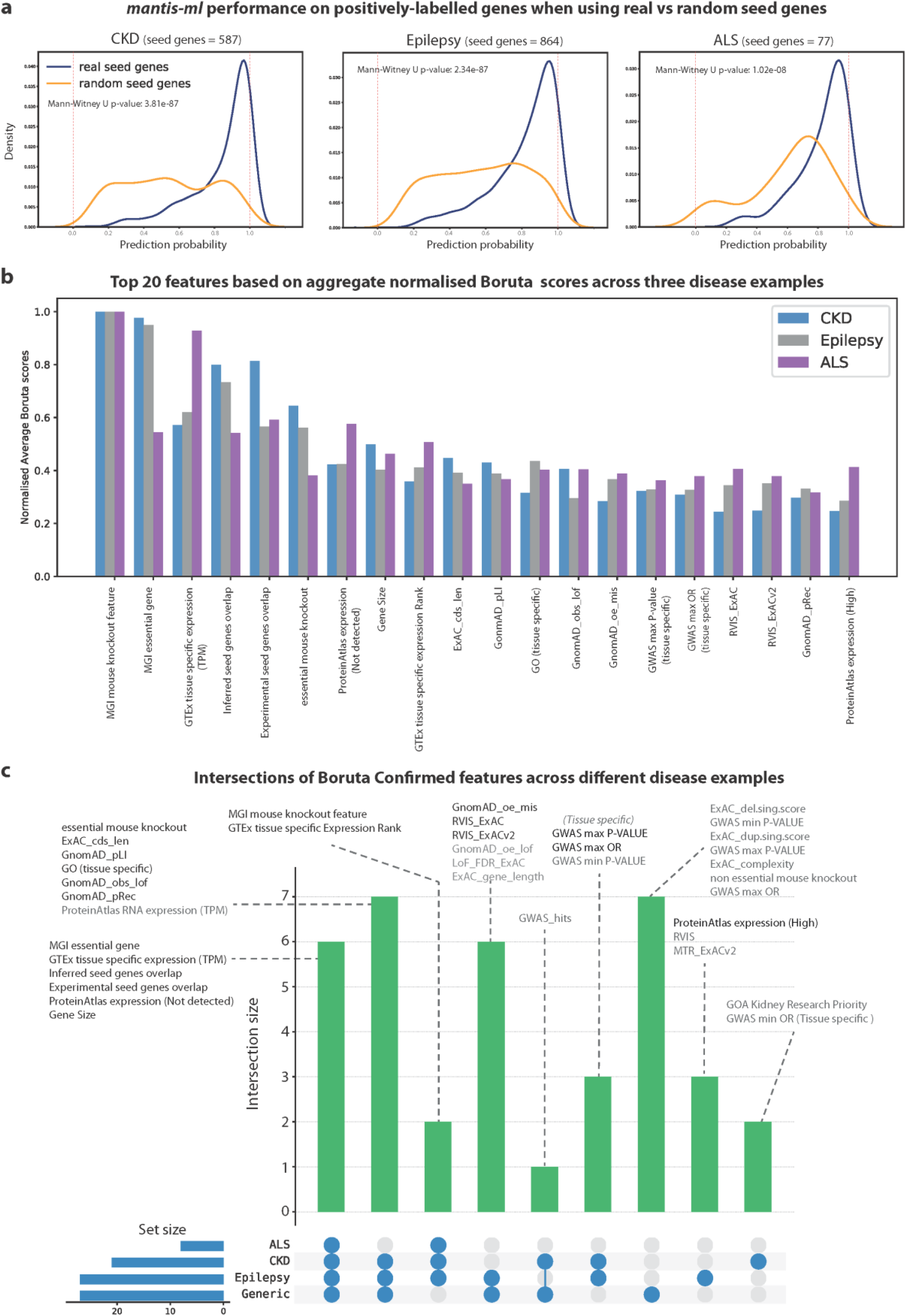
*Mantis-ml* performance sensitivity on seed genes and consensus of top feature contributors across different disease examples based on the Boruta algorithm. a) Prediction probability distributions from positively labelled (seed) genes across the three disease examples, when selected from Human Phenotype Ontology vs random assignment. Random seed genes are predicted with almost uniform probability distribution, while real seed genes successfully get ranked to the top of the spectrum (probability values close to 1). b) Top feature contributors per disease example based on sum of normalised average Z-scores returned by Boruta. Features are ranked merely based on their Z-scores without checking if they reach significance level in each disease case to be considered as ‘Confirmed’ features. c) Intersection of confirmed Boruta features across the three disease examples (CKD, Epilepsy and ALS) and the Generic disease classifier. Features with black font text correspond to the top-20 features shown above, based on the aggregate normalised Boruta scores.

### Mouse model phenotypes, tissue expression and protein-protein interactions recurrently among top feature contributors in *mantis-ml* predictions

Although *mantis-ml* focuses on identifying features highly predictive of known seed genes, we expect that some features are generally strong predictors agnostic to specific disease. *mantis-ml* integrates more than 1,200 features in total (**Fig. 1**) that are subset according to the disease under study. A typical disease example will be trained on around 100 features from *mantis-ml*, prior to any pre-processing. We sought to explore the contribution of each of the features during stochastic positive-unlabelled learning across all three examined disease cases. We employed the Boruta algorithm based on a Random Forest classifier, across 100 random balanced gene subsets with 10-fold cross-validation (see Methods). The Boruta algorithm provides an unbiased assessment of feature contribution as it constructs artefactual features (‘shadow’ features) from random permutations of each of the actual features of a dataset and then iteratively confirms or rejects the original features based on their Z-score distances from the importance levels achieved by the random (shadow) features. We have run Boruta for CKD, Epilepsy and ALS and extracted the consensus profile of feature importance in each disease case across 100 stochastic iterations (**Suppl. Fig 9a-c**; see Methods). We then normalised the Z-scores among the three disease cases (min-max normalisation) to compare the relative importance of each feature in the different diseases and provide a disease-agnostic consensus of feature importance profile (**Fig. 3b,c**).

The consensus of feature selection across CKD, Epilepsy and ALS reveals mouse model phenotypes and tissue-specific expression (Kidney, Brain and Brain/Skeletal Muscle, respectively, based on ProteinAtlas and GTEx) as consistently highly important contributors. Specifically, the ‘MGI mouse knockout feature’ is the top contributor in all three cases which reflects the high importance of tissue-specific animal models for validation of disease phenotypes in humans as well. This feature captures human genes with mouse orthologs that are associated with a ‘High-level Mammalian Phenotype’ (based on MGI) relevant to the disease under study (see Methods). Moreover, human orthologs of mouse genes that have been found to be essential for basic developmental functions and/or survival in both species (‘MGI essential gene’) are the next top contributors in CKD and Epilepsy and among the top contributors for ALS. Tissue-specific expression (‘GTEx tissue specific expression (TPM)’, ‘ProteinAtlas expression (Not detected)’ and ‘GTEx tissue specific expression Rank’) follow in the order of consistent feature importance, with a particularly high contribution to ALS, compared to the other two disease cases. A very interesting outcome of the Boruta algorithm is the emergence of protein-protein interactions-related features (‘Inferred seed genes overlap’, ‘Experimental seed genes overlap’) being ranked in the top five important features. We have constructed these features to capture the ratio of known (seed) genes interacting directly with each gene based on either ‘Experimental’ or ‘Inferred’ prediction from the *InWeb_IM* resource^8^ (see Methods). Finally, tissue-specific Gene Ontology (GO) terms, intolerance scores based on ExAC and GnomAD (‘GnomAD_pLI’, ‘GnomAD_obs_lof’, ‘GnomAD_oe_mis’, ‘GnomAD_pRec’ and ‘RVIS’), tissue-specific GWAS metrics (‘GWAS max P-value’, ‘GWAS max OR’) and gene size complete the picture of the most contributing features for classification of disease-associated genes in each of the examined disease examples.

### Application of *mantis-ml* predictions to triage results from exome-wide cohort association studies

The rapid development of next-generation sequencing (NGS) technologies in recent years has led to the ubiquitous application of large-scale genomic studies by large genomic and/or healthcare institutions for research and diagnostic purposes. Of special interest are large association studies that assess the enrichment of rare predicted deleterious variants in a collection of disease-ascertained cases compared to an available control population. Depending on the contribution of individual genes to disease risk these studies can provide experiment-wide significant results^17,18^, but more often they yield hundreds of highly-ranked genes of interest that do not exceed the genome-wide statistical significance thresholds given the available cohort size. Biologically relevant genes will reside among those highly ranked genes (lowest p-values but not experiment-wide significant); however, they will be surrounded by stochastic signals representing the natural trail of the null distribution. Teasing apart the biological signals from stochastic signals among the top ranks is a key challenge for most high-throughput genomics screens. The importance is that some of the gene candidates are then often pursued by downstream bioinformatic analysis often involving laborious post hoc manual or semi-automated search through existing literature and this process can be biased by a differing filtering process designed by each individual researcher based on their prior experience. Thus, different researchers adopting different resources will ultimately result in triaging different genes from each other.

*mantis-ml* eliminates subjectivity and post hoc design from gene prioritisation by using a standardised set of community knowledge and collectively assessing all interactions (both linear and non-linear) between multiple features that may characterise genes’ behaviour with respect to disease phenotypes. Here, we want to demonstrate the utility of leveraging the power of *mantis-ml* predictions to provide orthogonal information to support triaging the candidate genes found among results from Whole Exome Sequencing (WES)-based association studies. We selected one study per disease category where genes have been previously ranked based on the significant case-enrichment of various types of qualifying variants (e.g. putative loss-of-function [pLoF], missense, synonymous, etc.) between cases and controls for the respective cohort. Specifically, for CKD we are cross-referencing a study examining the preponderance of rare pLoF and other types of rare variants across a population of 3,150 cases and 9,563 controls^19^. Another study with 640 individuals with familial genetic generalised epilepsy and 3,877 controls has been employed for further triaging based on the *mantis-ml* predictions for the Epilepsy disease case^17^. Finally, for ALS we are using the results from a study looking into the collapsing analysis results of nearly 3,000 ALS patients compared against 6,405 controls^20^.

In each example, we sought to ask the question of whether the lowest p-values from the exome-wide association statistics (highest ranked from cohort studies) are significantly preferentially enriched for genes achieving *mantis-ml* highest predictions for that corresponding disease. To this end, we apply a hypergeometric enrichment test asking whether the top 5% of *mantis-ml* predictions in each disease case are preferentially overlapping with the top signals (lowest p-values) from the cohort-level association studies (i.e., the published rare-variant gene-based collapsing analysis gene lists). To strengthen our experimental design with negative control components, we assess this enrichment across several differing classes of qualifying variants including synonymous and permutation-based gene-ranks, which should represent the expected null to contrast the more biologically interesting models, such as the pLoF and ultra-rare deleterious missense collapsing analyses. To highlight whether there is significant enrichment among the lowest cohort-level association statistic p-values we are focusing our enrichment assessment of the top *mantis-ml* predictions among the top ranked statistically significant genes (P < 0.05) from the respective rare-variant gene-based association study. We observed that in all three disease examples, *mantis-ml* predictions overlap significantly with the exome study ranking for LoF variants (**Fig. 4a**). Notably, the significance of enrichment for LoF signal within *mantis-ml* predictions is the highest compared to all other ranked gene lists that are formed based on other type of qualifying variant classes that have been studied (synonymous, common, missense and shuffle/random permutation). Enrichment of high *mantis-ml* predictions (top 5% per disease) among LoF-associated genes (P-value < 0.05 from the published collapsing analysis results) is always statistically significantly different from a shuffled (randomised LoF-associated gene list) enrichment signal (Mann-Whitney U P-value=2.91×10^−300^, 5.53×10^−147^, 4.17×10^−194^ for CKD, Epilepsy and ALS, respectively). Moreover, enrichment of high *mantis-ml* predictions among top ranked LoF-associated genes is also always statistically significantly different from the enrichment signal from genes associated with synonymous variants (Mann-Whitney U P-value=1.34×10^−125^, 1.60×10^−33^, 2.36×10^−84^ for CKD, Epilepsy and ALS, respectively; considering ‘Dom_coding’ as the comparator class in ALS due to lack of a real synonymous collapsing analysis gene list in the published analysis).

**Fig. 4.**
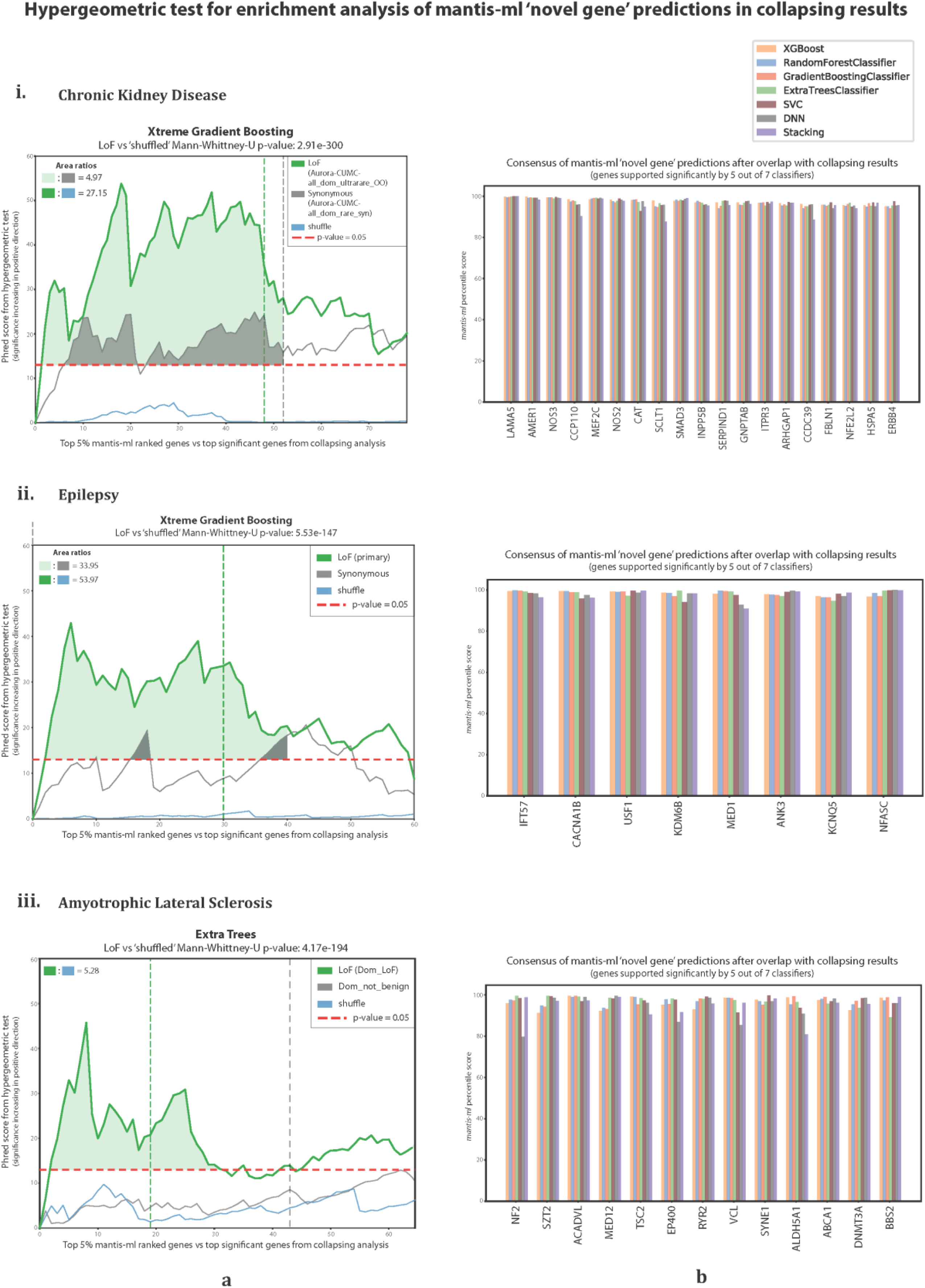
Cross-validation of *mantis-ml* predictions with cohort-level rare-variant association studies. a) Hypergeometric test enrichment of disease-specific *mantis-ml* predictions with collapsing analysis results from CKD (i), Epilepsy (ii) and ALS (iii) cohorts. The horizontal dashed red line corresponds to the significance threshold of p=0.05 for the hypergeometric tests. Where the plot(s) go above this line highlight significant enrichment of mantis-ml top gene predictions being enriched for among the population genomic collapsing analyses. The vertical dashed lines, coloured based on the different classes of qualifying variants, indicate where the last of top ranked genes from the collapsing analyses that achieved a p-value < 0.05. The highlighted areas (light green and grey) represent the magnitude of enrichment signal for Loss-of-Function and synonymous variants identified both from the collapsing analyses (p < 0.05) and the hypergeometric enrichment test against *mantis-ml* predictions (p < 0.05). b) Consensus of genes-of-highest-interest (novel) satisfying the significance threshold criteria in both the collapsing analysis results and the hypergeometric and supported by five out of seven classifiers used by *mantis-ml* in the CKD (i), Epilepsy (ii) and ALS (iii) disease examples.

We further quantified the enrichment signal by calculating the ratios of areas under curve between the LoF and synonymous collapsing analysis signals for CKD, Epilepsy and the total areas covered by the LoF enrichment signal for ALS, due to lack of a synonymous-associated signal in the respective published study. The synonymous enrichment signal serves as a negative control (technical baseline) since we expect genes prioritised during collapsing analysis based on synonymous variants not to be associated with pathogenicity, in general. We sought to explore how each of the seven classifiers performed when overlapped with top ranked WES-based gene lists. We observe that the best AUC performing classifiers per disease category in *mantis-ml* also ranked among the top three classifiers in terms of area under curve ratios and/or total LoF area (**Suppl. Fig. 10-12**). Other classifiers also perform comparably or slightly better, which is in concordance with the similar AUC performance achieved from differing classifiers retrieved from the original *mantis-ml* training, again, reinforcing the consistency and robustness of the framework irrespective of chosen classifier.

By applying the hypergeometric test for *mantis-ml* prediction enrichment in published case-control association studies, we can eventually extract a consensus list of predicted novel genes-of-highest-interest that satisfy both the hypergeometric test (p<0.05) and the collapsing analysis statistical significance threshold (p=0.05). We can further select the subset of genes that were supported by at least five of the seven classifiers in each disease case (**Fig. 4b**). Eventually, we highlight 19 (CKD), 8 (Epilepsy) and 13 (ALS) novel (unlabelled) genes-of-highest-interest extracted by overlapping the collapsing analysis results with the *mantis-ml* predictions across multiple classifiers. To validate this cross-validation approach we also assessed the results retrieved with regards to known (labelled) genes for each disease. For example, in CKD we observe that some of the most well-established CKD-associated genes (*PKD1*, *PKD2*, *COL4A1*, *COL4A3*, *COL4A4*, *COL4A5*) rank in the top 9 genes among the 17 known CKD genes that achieved both a collapsing analysis p<0.05 and hypergeometric test p<0.05 (**Suppl. Fig. 13**).

The unlabelled genes-of-highest-interest represent a collection of genes that were not among the HPO derived set of seed genes in the initial process of *mantis-ml*. Given the dependency on OMIM, some of these novel genes of highest interest (highly ranked by publish case-control studies and also by *mantis-ml* predictions) might have existing literature support for disease-association to act as a further positive control; however, with the caution that some might have been prioritised in literature on basis of similar underlying knowledge leveraged by *mantis-ml* to predict them as genes-of-interest. We looked for references in literature for the top suggested novel genes per disease by *mantis-ml* and have found supporting evidence for several genes. For instance, *LAMA5* has been reported in the last two years to be co-inherited with *COL4A5* in familial hematuria^21^ and hosting variants that may affect pediatric nephrotic syndrome^22,23^. Moreover, *NOS3* and *NOS2*—although not associated with CKD in OMIM —have already been implicated in Chronic Kidney Disease in a fair amount of studies^24,25,26,27^, while *MEF2C*, a gene typically associated with mental disorders, has also been associated with estimated Glomerular Filtration Rate (eGFR) or proteinuria^28^. *SCLT1* deficiency has been linked with cystic kidney^29^ and *SMAD* genes (including SMAD3) have been reported to affect CKD progression when dysregulated^30^. *INPP5B* impairment has been associated with severe renal phenotypes, such as proximal tubule endocytosis^31^, while targeting *NFE2L2* (NRF2) has been tested for prevention of Kidney Disease Progression^32^.

With regards to the top novel epilepsy-associated predictions, *CACNA1B* is associated with voltage-gated calcium channel which is subsequently implicated in epileptic phenotypes^33,34,35^. *USF1*-deficiency in mouse (in combination with *USF2* knockout) has been shown to cause epileptic seizures^36^, suggesting its important role for normal brain function. Furthermore, *KDM6B* is associated with neuronal survival^37^ and when haploinsufficient it causes severe seizures^38^. *ANK3* has also been reported to be involved in epilepsy^39,40^. Loss-of-Function and Gain-of-Function variants in *KCNQ5* have been shown to cause epileptic encephalopathy^41^.

As for the ALS consensus novel predictions, missense variants in *SYNE1* have been reported to be associated with a multisystemic neurological phenotypic spectrum that includes amyotrophic lateral sclerosis^42,43,44^. *ALDH5A1* is significantly down-regulated in the spinal cord of an ALS murine model^45^ while *ABCA1* is among the altered genes in frontal cortex of ALS samples^46^. Finally, motor neurons in human ALS show significant abnormalities in *DNMT3A*, which is also over-expressed in synapses in mice with motor neuron degeneration^47^.

### Visualisation of cross-validated *mantis-ml* predictions and downstream analysis

Visualising the consensus novel and known genes from the overlap of *mantis-ml* predictions with rare-variant collapsing analysis results is an important component to ensure appropriate interpretation and accessibility to the data, findings and interpretation. Principal Component Analysis (PCA) performed for each of the three disease examples used in this study achieves a slight segregation of positive and unlabelled genes, with the consensus novel and know gene predictions tending to differentiate the most from the rest of genes (**Suppl. Fig. 14**). However, PCA fails to capture a high ratio of the total variance to be explained by its first 2 or 3 components (the variance explained by the first 3 components in each disease case is on average about 21%). Since PCA is representing the original features as linear combinations of the projected Principal Components, its inability to identify patterns of high variability in the entire gene set implies that the associations between the various collections of features driving gene predisposition to disease are likely to be non-linear.

Thus, we then apply two popular dimensionality reduction techniques that can identify non-linear patterns in the original high-dimensional feature space for each disease example: t-Distributed Stochastic Neighbour Embedding (t-SNE) and Uniform Manifold Approximation and Projection (UMAP). We observe that both methods map most of the consensus novel genes-of-highest-interest in the neighbour of distinct clusters of known genes (**Suppl. Fig. 15**). Both techniques capture patterns in more localised regions of genes while UMAP may also retain elements from the global structure more efficiently than t-SNE. By contrasting the two projections we may identify clusters of genes that are more likely to be close (i.e. similar) with each other in absolute terms of distance (similarity). Finally, *mantis-ml* tool provides interactive visualisations of all three projections (PCA, t-SNE and UMAP) for further inspection of gene clusters as well as offers all extracted two-dimensional representations of the original data space that allow further extraction of clusters of genes e.g. using HDBSCAN for further downstream analysis.

### Generic *Mantis-ML* Score for gene disease likelihood

One of the key requirements for the above three disease-specific applications of *mantis-ml* is a sufficient collection of known OMIM disease-associated genes based on HPO term linkage. A rich collection of genes will not always be available for many diseases of interest. Therefore, we wanted to further explore the generation of disease-agnostic *mantis-ml* predictions. To achieve this, we trained *mantis-ml* using all current OMIM disease-associated genes (4,041 in total; see Methods) as seed genes to create a *Generic Mantis-ML Score* (GMS) that can be used as a general estimate of gene disease likelihood. Although it does not take full advantage of tissue- and disease-specific features, the GMS could be an opportunity to prioritise genes among disorders that currently have insufficient knowledge on causal genes.

We calculated the OMIM-based GMS using six different classifiers: Random Forest, Extra Trees, Gradient Boosting, XGBoost, SVC and DNN (here, we excluded Stacking due to its much longer training time compared to the other classifiers as well as the resulting slightly lower AUC performance). Gradient Boosting was the top performing classifier (average AUC=0.84) followed by Random Forest, XGBoost and Extra Trees with comparable AUC scores (**Supp. Fig. 16**). Similar to the disease-specific cases, in the generic disease *mantis-ml* results, we observe a high concordance between the predictions from all classifiers with regards to known disease genes among the out-of-bag test sets. Specifically, of the 4,041 known genes, around 1,300 (32.2%) were consistently identified by all classifiers (achieving probability > 0.5). Another 380 known genes (9.4%) were further identified by at least five of the six classifiers. With regards to novel disease-associated genes, again the largest group of predicted genes (n=600) was predicted by all classifiers, and ∼320 novels genes by at least five classifiers. In both cases, DNN had the highest number of known and novel predictions identified solely by a single classifier (280 and 680 predicted genes, respectively). We provide the average probability scores returned by Gradient Boosting as the default ranking for GMS along with the respective percentile score for each gene (**Suppl. File 2; Table S4**).

Furthermore, we ran again the Boruta algorithm for a single (*L=*1) stochastic iteration to identify the most important features that drive gene classification based on the entire OMIM disease annotation and compare it against the respective features (where applicable) extracted from the disease-specific cases. We observe that mouse model phenotypes (‘MGI essential gene’ and ‘essential mouse knockout’), protein expression (based on GTEx and Protein Atlas), protein-protein interaction features (‘Inferred seed genes overlap’ and ‘Experimental seed genes overlap’) are still the top feature contributors as in CKD, Epilepsy and ALS (**Fig. 5a**). Additionally, gene length, Gene Ontology features and intolerance scores (RVIS and the scores based on GnomAD) rank highly in the normalised average Boruta score scale.

**Fig. 5.**
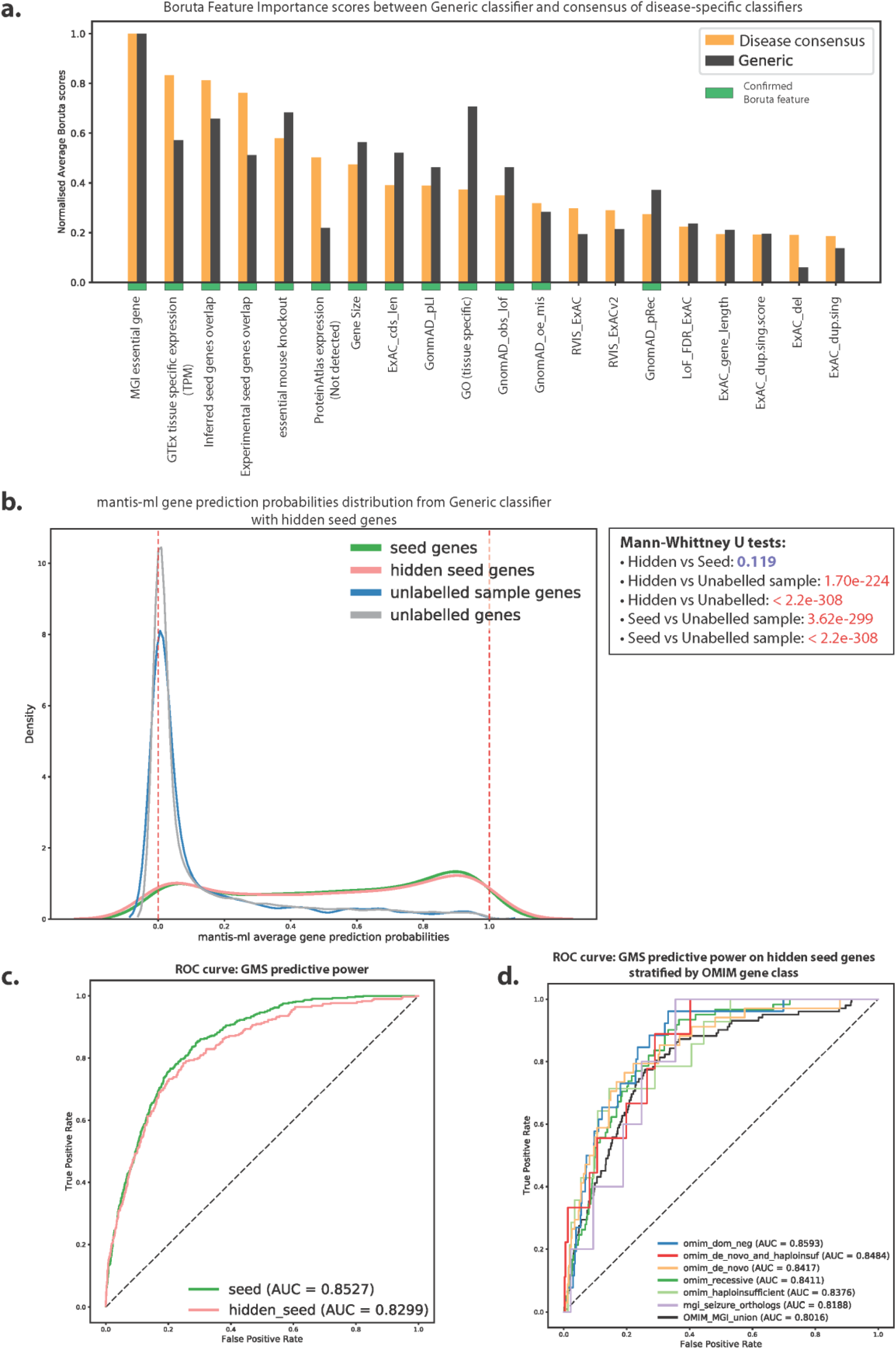
Generic disease *mantis-ml* classifier for estimation of gene disease likelihood. a) Comparison of consensus top feature contributors from CKD, Epilepsy and ALS with Generic *Mantis-ML* Score (GMS) feature importance scores. The consensus of disease-specific case is calculated as the mean of the normalised average Z-scores returned by Boruta for each disease case. b) Generic *mantis-ml* prediction probabilities across different gene classes. The ranking has been performed using 60% of the original seed genes set with the rest 40% of seed genes treated as unlabelled. The ‘unlabelled sample’ class includes a random sample from the unlabelled genes of equal size with the hidden genes set. Mann-Whitney U tests have been performed between the prediction probability distributions of all pairs of gene classes (p-values shown at the box on the right) to quantify their similarity degree. c) Predictive power of GMS to distinguish seed genes (unmasked and hidden) from unlabelled genes using a logistic regression classifier. d) Predictive power of GMS to distinguish different OMIM- and MGI-based hidden seed genes from unlabelled ones.

To validate the Generic disease classifier results, we explored the ability of *mantis-ml* to correctly identify a set of known genes which has not been provided as part of the original seed gene set. Thus, we masked a random selection of 40% of the 4,041 seed genes (considering them as unlabelled) and then trained the generic *mantis-ml* algorithm using Gradient Boosting as the standard classifier, for *L=5* stochastic iterations. Notably, masked OMIM disease genes were predicted as well as the unmasked seed genes with no significant difference (Mann-Whitney U P-value=0.119; **Fig. 5b**). We explored the predictive power of GMS in this case in terms of distinguishing seed genes from unlabelled ones and retrieved AUC scores of 0.853 and 0.83 for unmasked and masked (hidden) seed genes, respectively (**Fig. 5c**).

We then assessed the ability of GMS to efficiently stratify different OMIM- and MGI-based gene classes based on prediction probabilities assigned to hidden seed genes only, to avoid any over-prediction bias from the retained seed genes (unmasked). The gene classes that we used have been defined in a previous work^9^ and the intersection with the hidden genes used in this case are: OMIM dominant negative (130 genes), OMIM de novo and haploinsufficient (46 genes), OMIM de novo (168 genes), OMIM recessive (304 genes), OMIM haploinsufficient (70 genes) and MGI seizure orthologs (25 genes). We also compile the union of these gene classes as the ‘OMIM_MGI_union’ set (510 genes). We observe a high predictive power in this classification task for GMS with AUC scores ranging from 0.82-0.86 across the different OMIM/MGI gene sets and 0.80 of the union of these genes (**Fig. 5d**). These scores are considerably higher from similar assessments using other metrics of genic intolerance: gnomAD_pLI, gnomAD_mis_z, RVIS_ExACv2, ExAC_cnv.score and LoF_FDR_ExAC. The AUC performance of these scores in the task of OMIM/MGI vs non-OMIM/MGI classification (**Suppl. Fig. 21**) were in the range of: 0.58-0.85 (gnomAD_pLI), 0.47-0.79 (gnomAD_mis_z), 0.47-0.70 (ExAC_cnv.score), 0.50-0.68 (RVIS_ExACv2) and 0.50-0.72 (LoF_FDR_ExAC). The respective AUC scores for the classification of the union of OMIM/MGI gene sets vs non OMIM/MGI genes were in the range of 0.48-0.60 across all these metrics. It is, however, important to note that these component metrics are based on more specific data-types, whereas *mantis-ml* leverages the information from a wider and diverse collection of features. The ability of *mantis-ml* to correctly classify hidden seed genes underlines its power to subsequently correctly identify other unlabelled genes and provide a biologically meaningful ranking.

Finally, we explored how GMS would perform when overlapped with disease-specific rare-variant collapsing analyses results, similar to our assessment above using the disease-specific *mantis-ml* predictions (**Supp. Fig. 17**). We noticed that there is still significant enrichment of the LoF signal in all three cases but at a much lower level compared to the disease-specific cases (LoF/synonymous area ratios with GMS: 1.18, 0.81, 2.21 and respective scores with disease-specific classifiers: 4.97, 33.95, Inf/Not-defined, due to lack a synonymous class). This highlights the added value of leveraging the disease-specific classifiers, built on top of disease-specific features, to efficiently identify genes associated with a disease when such information (seed genes) is available. At the same time, the residual enrichment signal that can still be observed when using the Generic *mantis-ml* classifier predictions provides applied support about the utility of GMS to provide biologically relevant gene predictions when coupled with large case-control association statistics in the absence of disease-specific domain knowledge.

## Discussion

Present day, the genomics community is generating and analysing large volumes of exome and genome sequence data to better understand the genetic architecture of rare and common complex disorders. Here, we introduce a novel multi-dimensional machine learning framework, *mantis-ml*, to support the triaging of the result of those large-scale exome-wide genetic read-outs to further aide the prioritisation of novel disease genes. *mantis-ml* takes its name from the Greek word *’μάντης’* which means *’fortune teller’*. Given the demonstrable predictive utility of *mantis-ml*, particularly when generated in disease-specific predictions, the gene candidates triaged by *mantis-ml* in combination with rare-variant association studies from whole-exome sequencing data facilitate a standardised and objective prioritisation of genes for further functional validation in *in vitro* and *in vivo* models; either supplementing existing bioinformatic prioritisation pipelines or its own. As the power of cohort-level studies increases through access to larger cohort of sequenced individuals, we expect to be able to elucidate even more refined associations of novel genes implicated in complex disease phenotypes.

One key limitation of *mantis-ml*, as with most machine learning frameworks, is the dependency on existing patterns. As such, *mantis-ml* is most powerful in identifying new disease-associated genes that might cause disease through an existing understood mechanism. Disease genes representing an entirely novel disease mechanism might not be as highly prioritised. However, there might be opportunities to explore unsupervised approaches by clustering the results from t-SNE and UMAP (which are part of the *mantis-ml* processing workflow) and run pathway enrichment analysis to detect clusters of genes with a profile closely associated with the disease under study. Moreover, *mantis-ml* currently supports bespoke disease-specific features only for Chronic Kidney Disease and Cardiovascular disease (e.g. CKDdb, GOA, exSNP, etc.). These sets of features could be expanded for additional diseases using relevant data resources that might enable even more refined stratification of genes in other disease categories. *mantis-ml* also supports inclusion of additional public or user internal features to contribute to the underlying predictions.

Opportunities for future technical expansion of this approach include exploring integrating autoencoders for fully feature agnostic dimensionality reduction as well as applying Graph Convolutional Networks to better leverage information from protein-protein interaction networks.

Our multi-dimensional gene prioritisation framework has demonstrable value when used together with results from any gene-based burden or collapsing analysis and is agnostic to the disease cohort, provided there is already ample knowledge of disease-associated genes. In the absence of such known disease gene lists, we also provide a GMS that could be adopted. We propose use of *mantis-ml* as an objective, standardised, fully quantitative and automated gene-prioritisation tool for disease-specific or disease-agnostic studies. Additionally, we provide it as a complementary tool when assessing putative disease genes extracted from completely orthogonal genetic studies; thus, reducing the required time for triaging top gene candidates from what is often a weeks-to-months process down to just a couple of hours.

## Methods

### Data availability & pre-processing

#### - Generic Resources

##### ExAC

Exome Aggregation Consortium (ExAC) data are available at: http://exac.broadinstitute.org/downloads (last accessed on 06/03/2019). We integrate all data from CNV Counts and Intolerance Scores (‘*exac-final-cnv.gene.scores071316*’) and the ‘*GeneSize*’ feature from the Functional Gene Constraint Scores (‘*fordist_cleaned_exac_r03_march16_z_pli\_ rec_null_data.txt*’).

##### Essential mouse genes

We integrate data from Georgi et al. (2013) that contain annotation for human orthologs of mouse genes that have been found to be essential for basic developmental functions and/or survival in both species (available at: https://doi.org/10.1371/journal.pgen.1003484.s022, last accessed on 06/03/2019). Both genes that have been identified as essential or non-essential are recorded and used as features by *mantis-ml*.

##### Genic-intolerance scores

We integrate two types of genic-intolerance scores: Residual Variation Intolerance Score (RVIS) and Missense Tolerance Score (MTR). RVIS scores (applied to EVS, ExAC and ExAC v2) are publicly available at http://genic-intolerance.org while MTR scores are publicly available at http://mtr-viewer.mdhs.unimelb.edu.au (both last accessed on 06/03/2019).

##### GnomAD

Genome Aggregation Database (GnomAD) data are publicly available at https://gnomad.broadinstitute.org/downloads (release 2.1, last accessed on 06/03/2019). We integrate all gene constraint scores in the *mantis-ml* framework. We retain for each gene all associated constraint scores that correspond to the canonical transcript, choosing the longest one in case there are more than one canonical transcript annotated for a gene. Aforementioned ExAC is a subset of GnomAD; however, both versions of the associated constraint scores are adopted.

##### GWAS (used both in the generic and disease-specific models)

Genome Wide Association (GWAS) data are publicly available at: https://www.ebi.ac.uk/gwas/docs/file-downloads (last accessed on 06/03/2019). We integrate data from ‘All associations’ (v1.0.2). For the disease-specific model we select all entries that contain any of the ‘*seed_include_terms*’ and ‘*additional_include_terms*’ and do not contain any of the ‘*exclude_terms*’ from *config.yaml* in their ‘DISEASE/TRAIT’ field. For the generic model we include all entries. In both cases, however, we filter out any entry with a p-value over the genome-wide significance threshold (p-value threshold: 5×10^−8^). Then, both for the disease-specific and generic model, we assign a True boolean flag to every gene that has at least one GWAS hit for any of the query terms specified. We also record the total number of GWAS hits per gene as well as the min/max p-values and min/max Odds Ratios associated with each gene.

##### MGI (generic)

Mouse Genome Informatics (MGI) data are publicly available at: http://www.informatics.jax.org/downloads/reports/index.html (last accessed on 06/03/2019). We are integrating data from three files: Genotypes and Mammalian Phenotype Annotations for Marker Type Genes excluding conditional mutations (‘*MGI_GenePheno.rpt*’), Mouse/Human Orthology with Phenotype Annotations (‘*HMD_HumanPhenotype.rpt*’) and Mammalian Phenotype Vocabulary in OBO v1.2, tab-delimited and OWL Formats (‘*VOC_MammalianPhenotype.rpt*’). We combine all data from these files to link human with mouse orthologs and their associated high-level mammalian phenotype descriptions and IDs. Gene labelling for this feature is performed by string matching of the ‘*include_terms’* and ‘*additional_include_terms’* from the given *config.yaml* file with the ‘High-level Mammalian Phenotype ID’ field in *hmd_human_pheno.processed.rpt*. Finally, we also annotate all genes that are associated with a ‘Lethal’ phenotype with a True boolean flag. The MP IDs associated with a ‘Lethal’ phenotype are: 0002058, 0002080, 0002081, 0002082, 0002083, 0006204, 0006205, 0006206, 0006207, 0006208, 0008527, 0008569, 0008762, 0009850, 0010768, 0010769, 0010770, 0010831, 0010832, 0011083, 0011084, 0011085, 0011086, 0011087, 0011088, 0011089, 0011090, 0011091, 0011092, 0011093, 0011094, 0011095, 0011096, 0011097, 0011098, 0011099, 0011100, 0011101, 0011102, 0011103, 0011104, 0011105, 0011106, 0011107, 0011108, 0011109, 0011110, 0011111, 0011112, 0011400, 0013292, 0013293, 0013294.

#### - Resources filtered by tissue/disease

##### GTEx

Genotype-Tissue Expression (GTEx) data are publicly available at: https://gtexportal.org/home/datasets (V7, last accessed on 06/03/2019). We integrate RNA-Seq data that contain the median TPM expression values by tissue (‘*GTEx_Analysis_2016-01*-*15_v7_RNASeQCv1.1.8_gene_median_tpm.gct*’, last accessed on 06/03/2019). For the tissue-specific model case, we subset the GTEx tissues that match the strings defined in the ‘*tissue*’ and ‘*additional_tissues*’ fields in *config.yaml* and then aggregate all values by gene across all tissues.

Additionally, we assign a rank for each gene based on the aggregate expression across all matching tissues (ranks = {1, 2, 3,…} in order of decreasing expression). Genes with overall expression less than the median among all genes are assigned the same rank, equal to the total number of genes, to increase signal-to-noise ratio for the most highly-expressed genes. As for the disease-generic model case, we keep expression values across all tissues and retain them for each gene as separate features, while no rank is computed in that case.

##### Human Phenotype Ontology

The Human Phenotype Ontology (HPO) data are publicly available at: http://www.human-phenotype-ontology.org. We are using Build #154 from HPO to annotate disease-associated genes (last accessed on 04/03/2019). We are using by default the ‘*ALL_SOURCES_FREQUENT_FEATURES_\ genes_to_phenotype.txt*’ file (provided by the HPO consortium), as our reference annotation file to exclude phenotypic features that are observed occasionally (present in 5–29% of the cases), rarely (present in 1–4% of the cases) or not at all (present in 0% of the cases). Annotation is performed by selecting genes whose ‘*HPO-Term-Name*’ contains any of the ‘*seed_include_terms*’ and does not contain any of the ‘*exclude_terms*’ from the *config.yaml* configuration file.

##### Human Protein Atlas

Human Protein Atlas data are publicly available at: https://www.proteinatlas.org/about/download (version 18.1, last accessed on 06/03/2019). We integrate two types of data from Human Protein Atlas: Normal tissue data (*normal_tissue.tsv*), which contain levels of expression for each gene in different tissues and cell types (categorical variable: ‘Not detected’, ‘Low’, ‘Medium’, ‘High’) and RNA gene data (*rna_tissue.tsv*), which contain TPM expression values for each gene by Sample (where ‘*Sample*’ in this case is similar with the ‘*Tissue*’ field from Normal tissue data).

We initially filter out all entries that have an ‘Uncertain’ value in the ‘Reliability’ field. For the disease-specific model case, we select all genes which contain any of the strings from ‘*tissue*’, ‘*seed_include_terms*’ and ‘*additional_include_terms*’ and do not contain any of the ‘*exclude_terms*’ from *config.yaml* file in their Tissue/Sample fields for Normal tissue and RNA gene data, respectively. Normal tissue data contain in general multiple values (levels of expression) per gene for the different cell types under each tissue type. We collapse all values for each gene by selecting the highest level found in a cell type within each tissue (‘Not detected’ < ‘Low’ < ‘Medium’ < ‘High’). With regards to RNA gene data, we aggregate all TPM values for each gene.

As for the generic disease model, expression levels from Normal tissue data are retrieved across all tissues, the highest level is retained for each gene and eventually we convert the four original levels into two: ‘Not detected’ and ‘Low’ are both considered as ‘Low’ and ‘Medium’ and ‘High’ are both considered as ‘High’. This transformation is performed to increase signal-to-noise ratio on this feature when looking at expression across all tissues. Finally, RNA gene data are aggregated by gene for each Sample.

##### InWeb_IM

InWeb_IM data (human protein-protein interaction network data, Li et al. 2017) are publicly available at: https://www.intomics.com/inbio/map.html#downloads (‘*inBio_Map_core_2016_09_12.zip*’, last accessed on 06/03/2019). Protein-protein interactions are characterised as ‘inferred’ or ‘experimental’ based on the validation degree recorded for each interaction in the original analysis. We untangle all interacting genes for each gene by validation type (‘inferred’ or ‘experimental’) and during analysis we record the ratio of interacting genes that belong to the seed genes (positively labelled genes) in each disease-specific run.

##### MGI

Data compilation is performed as described at the ‘MGI’ section in ‘Generic Resources’. For the disease-specific model, annotation is performed by selecting all genes whose linked phenotypes contain any of the strings from ‘*seed_include_terms*’ and ‘*additional_include_terms*’ and do not contain any of the ‘*exclude_terms*’ from *config.yaml*.

##### MSigDB

Molecular Signatures Database (MSigDB) data are publicly available at: http://software.broadinstitute.org/gsea/downloads.jsp (v6.2, last accessed on 06/03/2019). We integrate data from the c5 gene set (gene ontology sets). For the disease-specific model case, we select all gene ontology terms that contain any of the strings from ‘*tissue*’, ‘*seed_include_terms*’ and ‘*additional_include_terms*’ and do not contain any of the ‘*exclude_terms*’ from *config.yaml*. As for the disease-generic model, we retain all gene ontology terms. In both cases, gene ontology terms with less than 150 associated genes (0.08% of all genes) are filtered out to reduce the number of features with near-zero variance. In the current dataset this leaves 1,009 of 5,917 gene ontology terms.

##### OMIM

Online Mendelian Inheritance in Man (OMIM) data are available under licensing at: https://www.omim.org (‘*genemap2.txt*’, last accessed on 06/03/2019). We have restricted OMIM data to the subset of entries where the field ‘*Phenotypes*’ contains ‘*(3)*’, which reflects entries where the ‘*molecular basis for the disorder is known; a mutation has been found in the gene*’. OMIM annotation data are used only for extracting a disease-generic gene ranking. By default, all genes which contain ‘*(3)*’ in their ‘*Phenotypes*’ field are annotated as disease-associated genes (value ‘*All*’ in ‘*generic_classifier*’ parameter in the *config.yaml* file). Additional filtered layers of positive gene data are available by specifying different values for the ‘*generic_classifier*’ in *config.yaml*: a) ‘*AD*’ for selecting only genes that include ‘*Autosomal dominant*’ annotation in their ‘*Phenotypes*’ field, b) ‘*AR*’ for selecting only genes that include ‘*Autosomal recessive*’ annotation in their ‘*Phenotypes*’ field, c) ‘*AD_only*’ for selecting only genes that include ‘*Autosomal dominant*’ annotation and at the same time do not contain ‘*Autosomal recessive’* annotation in their ‘Phenotypes’ field, d)) ‘*AR_only*’ for selecting only genes that include ‘*Autosomal recessive’* annotation and at the same time do not contain ‘*Autosomal dominant’* annotation in their ‘*Phenotypes*’ field.

All string-matching operations are case insensitive.

#### - Disease-specific Resources (currently supported)

##### i. Chronic Kidney Disease (CKD)

###### CKDdb

Data from the Chronic Kidney Disease database (CKDdb) are available at: http://www.padb.org/ckddb (last accessed on 08/03/2019). We annotate each gene that has been associated with a renal disease with a True boolean flag and also record the total number of studies in CKDdb that support this evidence.

###### nephQTL

eQTL data for the glomerular and tubulointerstitial tissues (NephQTL) are available at: http://nephqtl.org. This database contains *cis*-eQTLs of the glomerular and tubulointerstitial tissues of the kidney found in 187 participants in the NEPTUNE cohort. For each gene and tissue, we record the expected number of eQTLs, the probability of not having eQTLs and the False Discovery Rate (FDR).

##### ii. Cardiovascular Disease

###### exSNP

Data from the database of expression associated SNPs (exSNP) are available at: http://www.exsnp.org/Download (last accessed on 08/03/2019). We are integrating disease associated high confidence (r^2^ > 0.8) eQTLs for Coronary Artery Disease and Hypertension and record the total number of eQTLs associated for each gene in each condition.

###### Adipose eQTLs

Data for adipose eQTLs identified at GWAS loci for cardiometabolic diseases and traits were retrieved from Civelek et. al, 2017 (Table S8). We record for each gene the total number of GWAS loci and cis eQTLs that have been associated for cardiometabolic traits.

###### Platelet eQTLs

Data for platelet eQTLs were retrieved from Simon et. al, 2016 (Table S2). Platelets have been shown to contribute to ischemic cardiovascular events^48^. We record for each gene the total number of heterozygous coding sites with a marginal eQTL effect (p < 10^−4^) and with 10 or more reads.

### Feature pre-processing

*mantis-ml* performs automatic feature pre-processing which includes filtering highly correlated features (parameter *eda_parameters* -> *high_corr_thres* in config,yaml; default value: 0.8). Additionally, features with a number of missing data over a certain threshold are discarded (parameter *eda_parameters* -> *missing_data_thres* in config,yaml; default value: 0.25). The remaining features with missing data ratio below the cut-off threshold are imputed with either a zero value or the median of the respective feature. Imputation with zero is performed either due to most of the genes representing a binary flag (‘non-existent’ or 0 for missing data, e.g. ‘MGI_mouse_knockout_feature’, ‘GOA_Kidney_Research_Priority’, etc.) or having extracted these features from computational or experimental studies that retrieve a biologically-relevant signal only from a specific set of genes, associated with the hypothesis under examination (e.g. ‘platelets_eQTL’, ‘adipose_GWAS_locus’ features, etc.). The features that are imputed with a median value are all GWAS and ExAC CNV-associated features, RVIS, MTR and gene-length. The selection of the median value for imputation in that case is due to different studies and/or resources being based on different global reference sets of genes, which requires extrapolation of these features to genes with missing values, since penalising them with a zero value would most likely not be representative of the actual gene behaviour with respect to these features. Finally, features are standardised to have mean zero value and unit variance.

### Stochastic positive-unlabelled prediction with standard classifier

The input data for *mantis-ml* are all coding genes, labelled based on known/unknown annotation for a disease, and accompanied with a large set of gene-level features extracted from public databases. To explore the potential of correctly classifying disease-associated genes, we initially set up a controlled environment with a single balanced dataset of positive and unlabelled data points from the CKD disease example. This dataset included over 1,000 genes (with positive:unlabelled ratio = 1:1.5). Within this balanced dataset we ran 10-fold Cross-Validation to assess *mantis-ml*’s predictive power in the out-of-bag dataset. Training was performed with seven different classifiers to assess their relative performance (Random Forests, Extra Trees, Gradient Boosting, Extreme Gradient Boosting, Support Vector Classifier, Deep Neural Net and Ensemble Stacking method consisted of a combination of these base classifiers). Fine-tuning of each of the classifiers was performed using Grid-Search with 10-fold Cross-Validation on a single random balanced dataset from CKD. We then streamlined this process across 10 random balanced datasets to perform a moderate size benchmarking test. Performance across all classifiers was comparable leading us to the choice to keep all classifiers as an option to be used by *mantis-ml* in each run. Eventually, we scaled up the positive-unlabelled learning task to the entire gene space, which is covered by a random partitioning of the unlabelled genes in combination with a random subset of the positive (seed) genes each time. The final ranking is extracted by averaging the prediction probabilities assigned to each gene from all the generated out-of-bag sets. This approach allows genes to ‘compete’ in a stochastic semi-supervised manner with each other and perform self-sorting considering that their respective features can capture enough variance of truly disease-relevant characteristics.

### Optimal classifier parameters selection with Grid Search Cross-Validation

Fine tuning for all *scikit-learn* based classifiers (Random Forest, Extra Trees, Gradient Boosintg, SVC) and for XGBoost was performed using GridSearchCV from *scikit-learn’s model_selection* module, over a pre-defined finite parameter grid space and tested on a single random balanced dataset from the CKD disease example. With regards to *keras*-based Deep Neural Networks, we developed a module that performs Grid Search with cross-validation (*dnn_grid_search_cv.py)*, including tuning of parameters such as size and number of hidden layers. This module currently supports simultaneous fine-tuning of up to 2 features but can otherwise fine-tune any DNN-related parameters in a single run, with sequential steps of optimisation that progressively select near-optimal features in a heuristic manner. Below are given the optimal features returned from each Grid Search with Cross-Validation run per classifier.

### Optimal selection of number *L* stochastic iterations in PU-learning

We examined the number of known and novel genes predicted for different numbers of stochastic iterations of PU learning using an Extra Trees classifier on a disease-specific example (**Suppl. Fig. 19**). We observed that the number of predicted known genes is practically insensitive to the number of stochastic iterations. However, the number of novel genes requires a certain number of iterations until it reaches a stable state, which is around 1000 genes. Specifically, novel genes count enters an oscillation zone around the stable state after *L*=50 iterations which is further stabilised after *L*=100 iterations. Eventually, we choose *L*=100 iterations as the default value for use with all PU-learning tasks with *mantis-ml* to increase the confidence level of gene probabilistic predictions.

In a more rigorous testing, we assessed the correlation of *mantis-ml* average prediction probabilities when run for different number of stochastic iterations. Ideally, a robust algorithm should capture the same average profile for each gene irrespective to the number of iterations *mantis-ml* has been trained on. Specifically, we ran *mantis-ml* for the following numbers of stochastic iterations: 1, 10, 30, 50, 70, 100, 150, 200. Pearson’s r correlation of the average *mantis-ml* prediction probabilities extracted from just one iteration compared to all other numbers of iterations is always > 0.984 (p<2.2×10^−308^). For any other pair of stochastic iterations > 1, Pearson’s correlations are in the range of 0.9976-0.999 (p<2.2×10^−308^). These predictions demonstrate the robustness of *mantis-ml* predictions irrespective to the number of stochastic iterations used. Thus, we suggest using *L*=10 iterations for an exploratory run on a disease-specific case and only run for *L*=100 in production to retrieve the final slightly more refined results.

### Dictionary of inclusion/exclusion query terms for the disease examples studied

**Table 1.** Optimal parameters for each classifier calculated with Grid Search and 10-fold Cross-Validation.

| <u>Classifier</u> | <u>Parameters selected with Grid Search</u> |
| --- | --- |
| **Deep Neural Network (DNN)** | • **hidden layers:** 2<br>• **nodes per layer:** [32, 32]<br>• **dropout ratio:** 0.3<br>• **L2 regularisation parameter:** 0.01<br>• **optimizer:** 'Adagrad'<br>• **epochs:** 50<br>• **batch_size:** 128 |
| **Extreme Gradient Boosting (XGBoost)** | • **learning_rate:** 0.01<br>• **n_estimators:** 300<br>• **max_depth:** 5 |
|  | • **min_child_weight:** 3<br>• **gamma:** 0<br>• **subsample:** 0.8<br>• **colsample_bytree:** 0.8<br>• **objective:** 'binary:logistic'<br>• **scale_pos_weight:** 1 |
| **Extra Trees** | • **n_estimators:** 100<br>• **max_features:** 'auto'<br>• **max_depth:** 15<br>• **min_samples_leaf:** 2<br>• **min_samples_split:** 5 |
| **Gradient Boosting** | • **n_estimators:** 500<br>• **max_features:** 'sqrt'<br>• **max_depth:** 20<br>• **min_samples_leaf:** 4<br>• **min_samples_split:** 5 |
| **Random Forest** | • **n_estimators:** 100<br>• **max_features:** 'auto'<br>• **max_depth:** 15<br>• **min_samples_leaf:** 2<br>• **min_samples_split:** 4<br>• **warm_start:** False |
| **Support Vector Classifier (SVC)** | • **C:** 0.01<br>• **kernel:** 'linear'<br>• **gamma:** 'auto'<br>• **probability:** True<br>• **shrinking:** True |

**Table 2.**
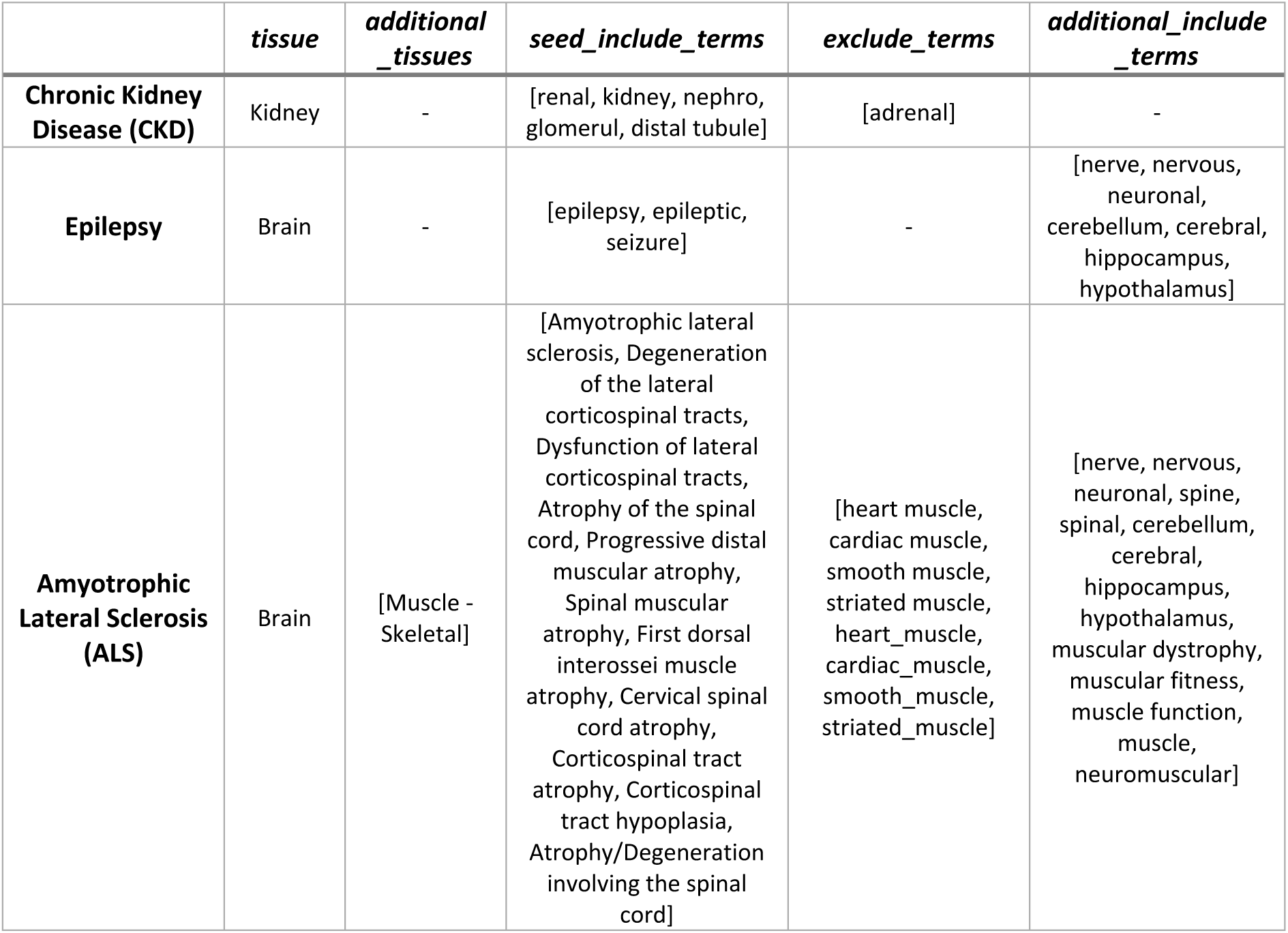
Query terms per disease category used in the configuration file for *mantis-ml* (*config.yaml*). All query terms are case insensitive and follow regular rules of wild card pattern matching. Cells with ‘-‘ should be given an empty list ‘[]’ as their value in the *config.yaml* file.

### Feature importance extraction with the Boruta algorithm

The Boruta algorithm has been ran on top of a Random Forest classifier trained on 100 random balanced datasets with 10-fold cross-validation and ran internally for 100 iterations. Boruta is assessing the importance of each feature by comparing its contribution with the one from random permuted features and eventually provides Z-scores that quantify the distance from that comparison. Upon each Boruta training cycle on a random balanced dataset, features are characterised as ‘Confirmed’, ‘Tentative’ or ‘Rejected’ and the full distribution of Z-scores is provided for each of them. Due to the stochastic nature of the positive-unlabelled learning implemented by *mantis-ml*, features may be characterised by different labels in different runs of the Boruta algorithm. We have thus defined a decision threshold to classify each feature as ‘Confirmed’, ‘Tentative’ or ‘Rejected’ based on its extracted labels across all Boruta runs. Specifically, for CKD and Epilepsy, features are eventually classified as ‘Confirmed’ when receiving this label across at least 90% of all Boruta runs. The decision threshold for the ‘Confirmed’ features classification has been set to 60% for ALS to compensate for the higher variance of extracted feature importance labels across all iterations, most likely due to the smaller set of seed genes in that case. In all three cases, features labelled as ‘Rejected’ in at least 90% of the cases are eventually classified as such, while the remaining features are characterised as ‘Tentative’ on the consensus labelling.

We have also added an option for *mantis-ml* to be trained using only the set of ‘Confirmed’ features extracted by the Boruta algorithm (parameter ‘*supervised_filters -> feature_selection: boruta’* in *config.yaml*). We then tested *mantis-ml*’s performance with each of the standard classifiers, separately, when using different configuration of features: using only Boruta confirmed features or using all features that survived after the feature pre-processing step (**Suppl. Fig. 18**). Training and prediction were performed using a set of 15 random balanced datasets from the CKD disease example. *mantis-ml* performed slightly but non-significantly better when using the entire feature set (average AUC: 0.817 vs 0.815; two sample t-test: P=0.515). However, since Boruta does not need to be run by default as part of the *mantis-ml* workflow, the default configuration retains the entire processed feature space and allows the user for further exploration by explicitly specifying ‘boruta’ as the feature selection algorithm in ‘*supervised_filters*’ field in *config.yaml*. We also tested the performance of Stacking classifier when fed with only the base classifier predictions vs employing the original feature space on top of the extracted base classifier predictions. Stacking classifier’s performance was considerably better when retaining the original feature space at the second layer of the ensemble training/prediction and this is the default configuration used by *mantis-ml*.

### *mantis-ml* package structure and code availability

Our *mantis-ml* framework has been built using Python on top of the *sckit-learn* and *keras* libraries. We have also employed the *Boruta* R package for feature selection based on the Boruta algorithm. The main components of *mantis-ml* are the *pre_processing*, *unsupervised_learn*, *supervised_learn*, *post_processing* and *validation* modules. The *pre_processing* module implements the functionality for compilation of the input feature table, which contains three classes of features: generic features (tissue/disease-agnostic), filtered by tissue and disease-specific features. Compilation of tissue/disease-specific features is performed using a curated dictionary of relevant query terms. Following data compilation, the *pre_processing* module implements the rest of its main functionality around feature pre-processing, exploratory data analysis and visualisation of features distribution. The *unsupervised_learn* performs dimensionality reduction on the processed feature set for visualisation purposes and extraction of 2-dimensional representations of the data for downstream analysis, such as clustering and pathway enrichment analysis. Processed feature tables are then passed on to the *supervised_learn* module for feature selection with Boruta and the stochastic positive-unlabelled learning task, which is the core of the *mantis-ml* workflow. Prediction probabilities extracted from this step are fed to the *post_processing* module for results aggregation and optionally overlapping with third-party studies (e.g. rare-variant cohort studies or any independently-generated ranked gene list) using the *validation* module.

The *mantis-ml* tool along with instructions to run can be found at the *mantis-ml-release* GitHub repository: https://github.com/astrazeneca-cgr-publications/mantis-ml-release

## Supplementary Data

**Supplementary File 1.** List of features used for disease-agnostic gene prioritisation in the *mantis-ml* machine learning framework.

**Supplementary File 2.** Tables with *mantis-ml* gene prioritisation scores (both raw prediction probabilities and percentile scores) for Chronic Kidney Disease, Epilepsy, Amyotrophic Lateral Sclerosis and the Generic *Mantis-ML* Score (GMS).

## Footnotes

This work was supported by funding from AstraZeneca.

---

**Suppl. Fig. 1.**
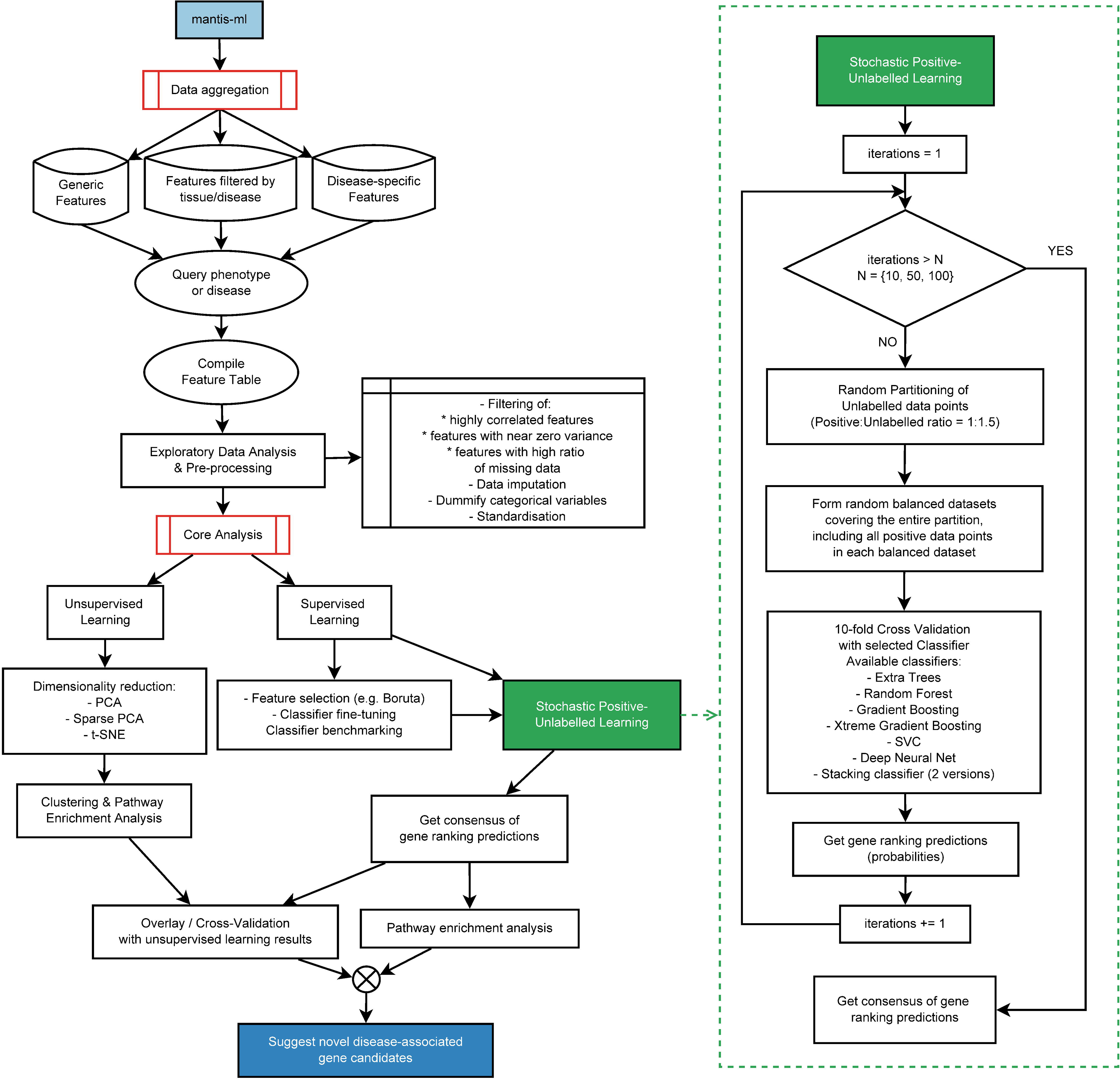
mantis-ml algorithm flowchart.

**Suppl. Fig. 2.**
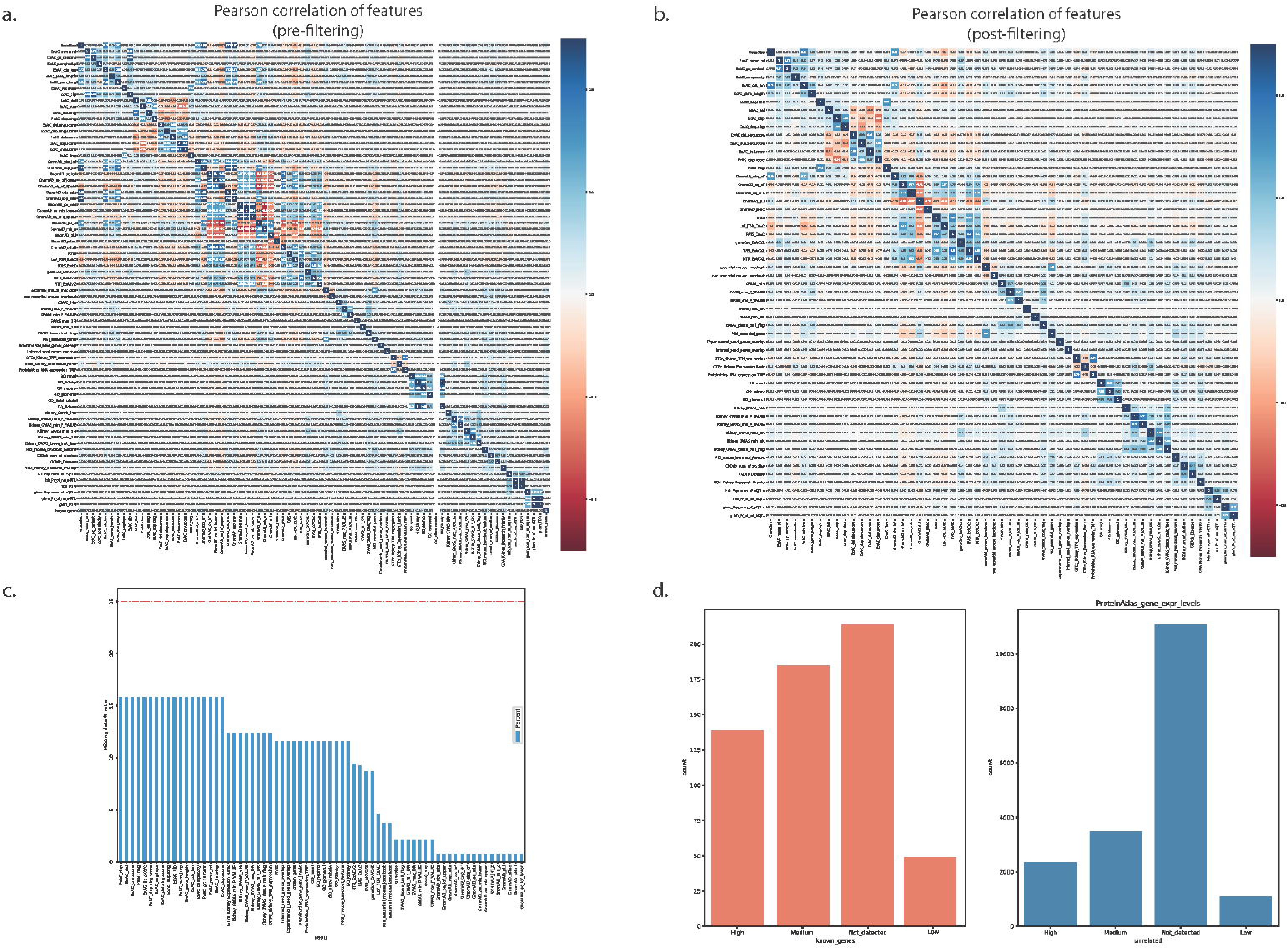
Data pre-processing and exploratory data analysis in Chronic Kidney Disease case: a) Pairwise feature correlation **before** filtering of highly-correlated features, b) Pairwise feature correlation **after** filtering of highly-correlated features, c) Missing data ratios and d) distribution of categorical feature profiles in positive and unlabelled genes.

**Suppl. Fig 3.**
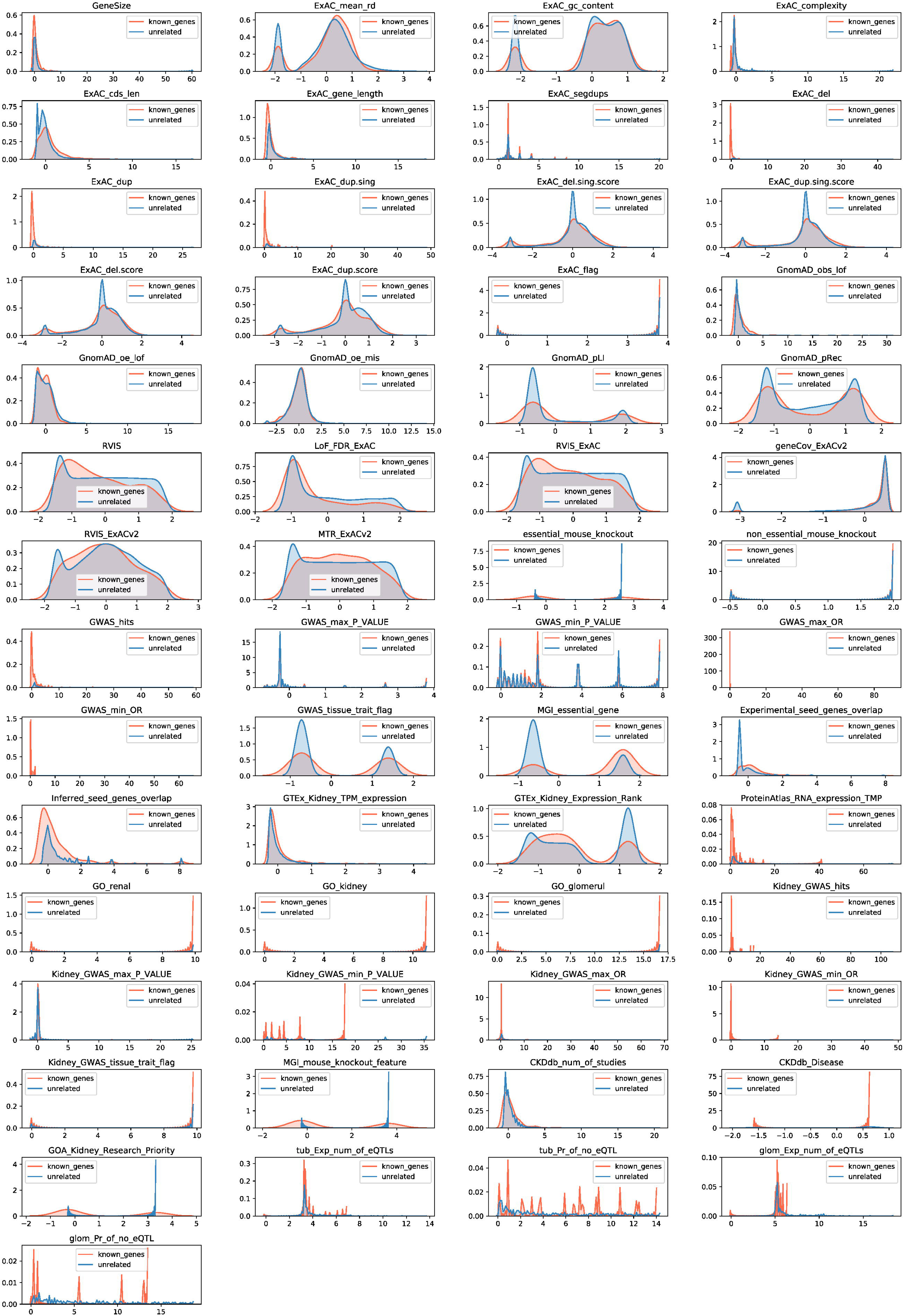
Exploratory data analysis in Chronic Kidney Disease case: distribution of numerical feature profiles in positive (known) and unlabelled genes

**Suppl. Fig. 4.**
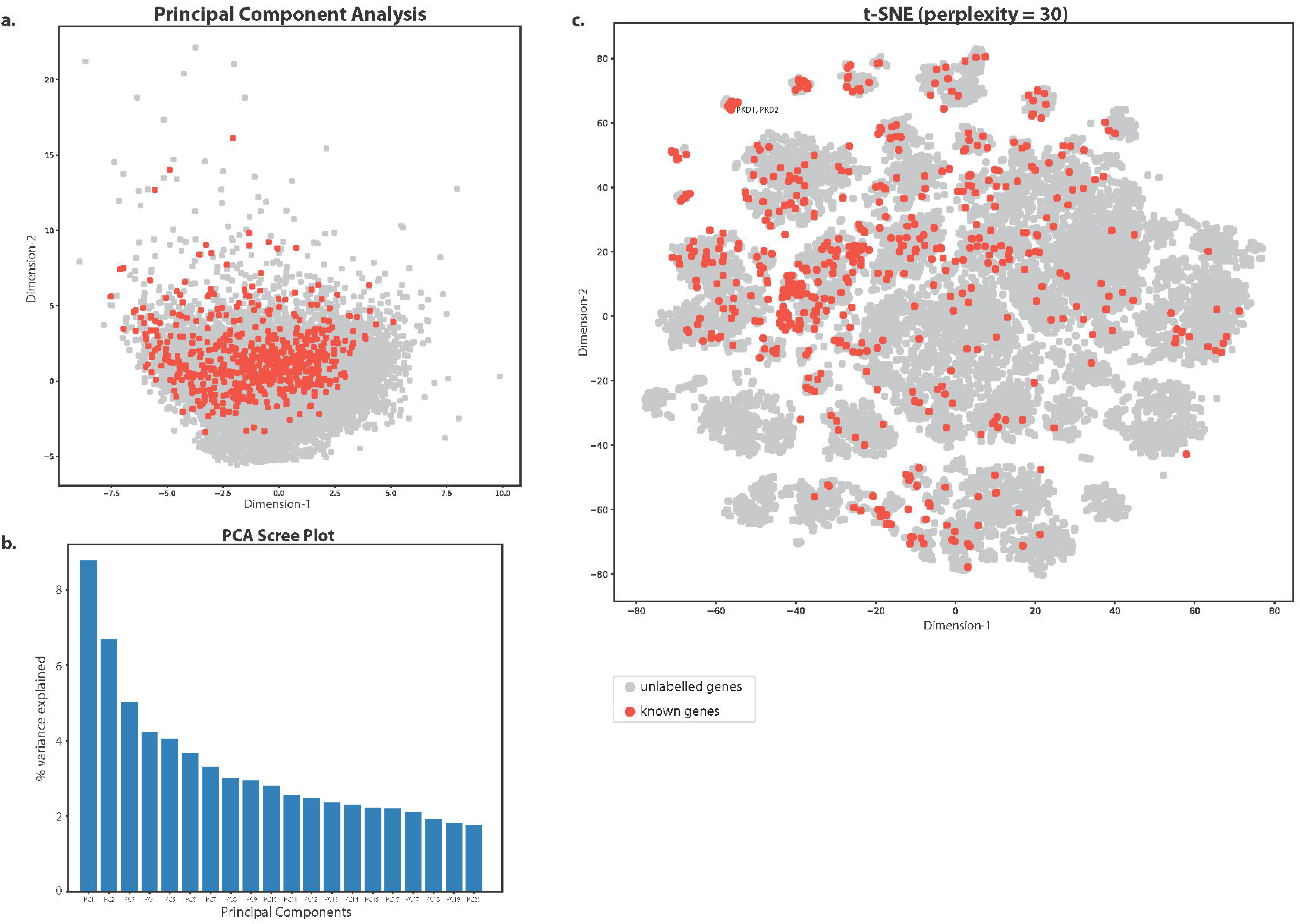
Dimensionality reduction on the Chronic Kidney Disease feature set: a) Principal Component Analysis, b) Scree plot from PCA with variancexplained by each of the calculated principal components and c) t-distributed Stochastic Neighbouring Embedding (t-SNE) using perplexity=30.

**Suppl. Fig 5.**
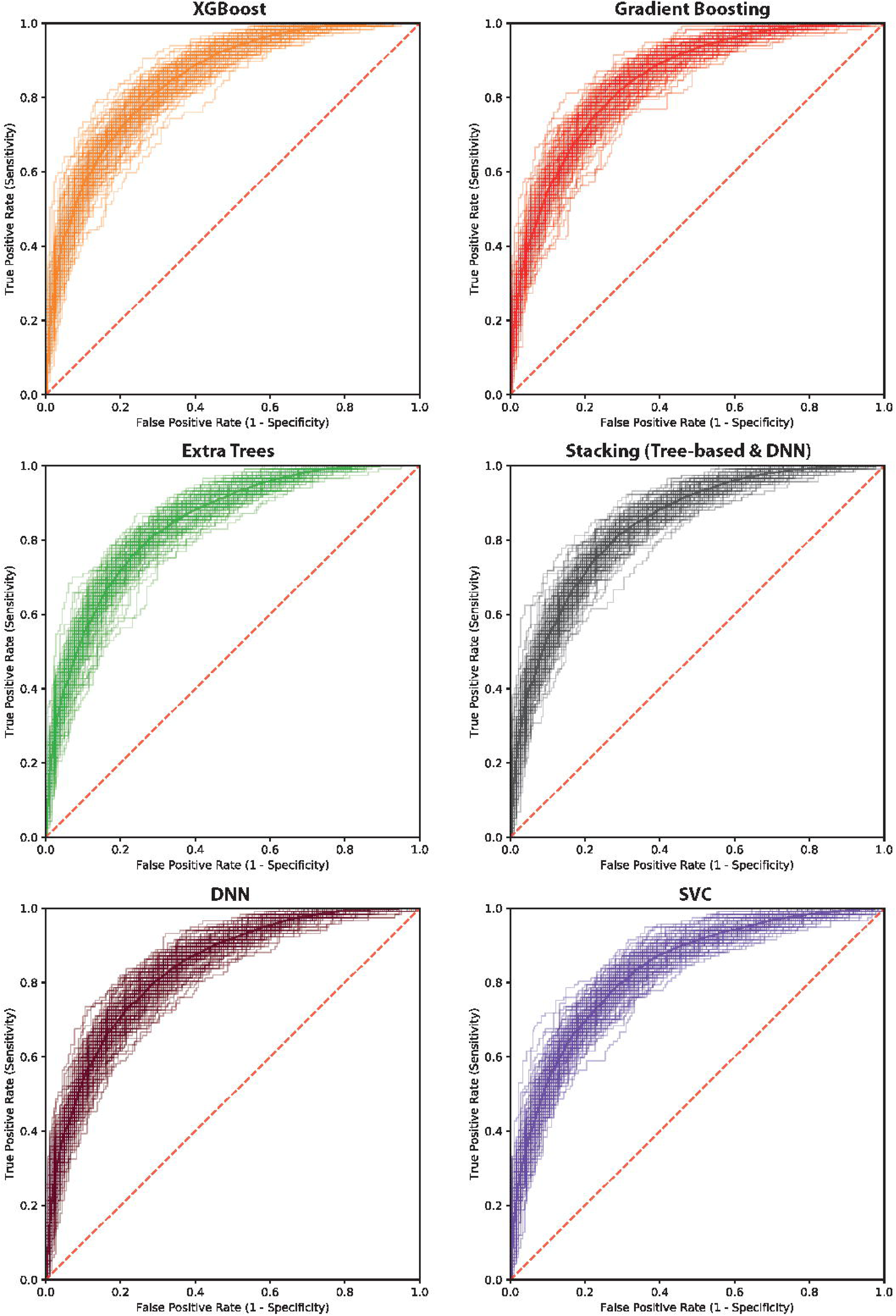
ROC curves from 10 batches of 10-fold Cross Validation with 6 different classifiers, in decreasing order of mean AUC: a) Random Forest, b) Xtreme Gradient Boosting, c) Gradient Boosting, d) Extra Trees, e) Stacking Classifier (1 st layer: Extra Trees + Random Forest + Gradient Boosting + SVC; 2nd layer: DNN), f) Deep Neural Net (2-hidden layers) and g) Support Vector Classifier.

**Suppl. Fig. 6.**
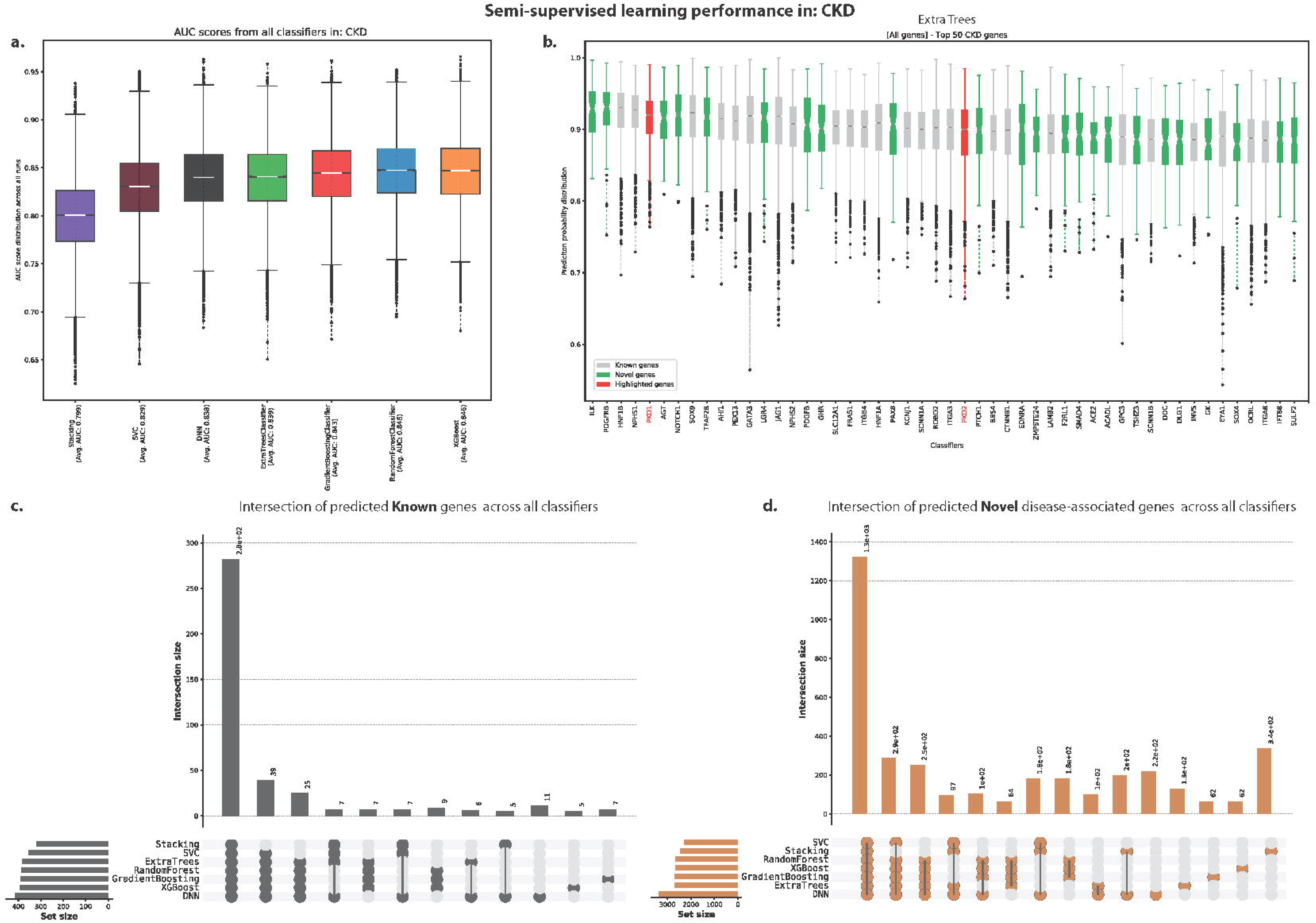
Mantis-ml performance on the Chronic Kidney Disease case, a) AUC score distribution per standard classifier used during mantis-ml training, b) Prediction probabilities from the top 50 (known and novel) genes predicted with Extra Trees as the standard classifier, c/d) Intersection sets of predicted known/novel genes across all classifiers.

**Suppl. Fig. 7.**
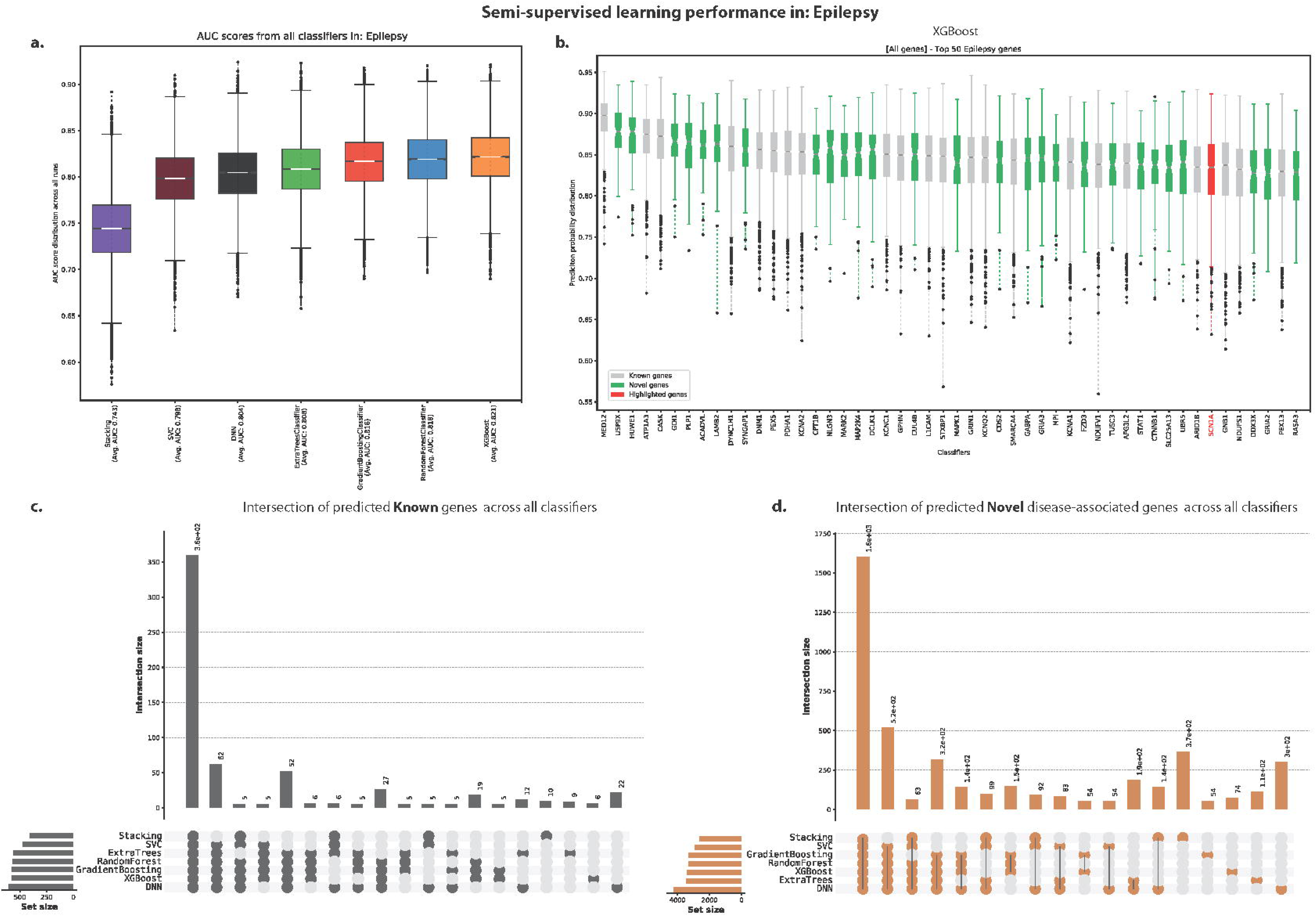
Mantis-ml performance on the Epilpesy disease case, a) AUC score distribution per standard classifier used during mantis-ml training, b) Prediction probabilities from the top 50 (known and novel) genes predicted with XGBoost as the standard classifier, c/d) Intersection sets of predicted known/novel genes across all classifiers.

**Suppl. Fig. 8.**
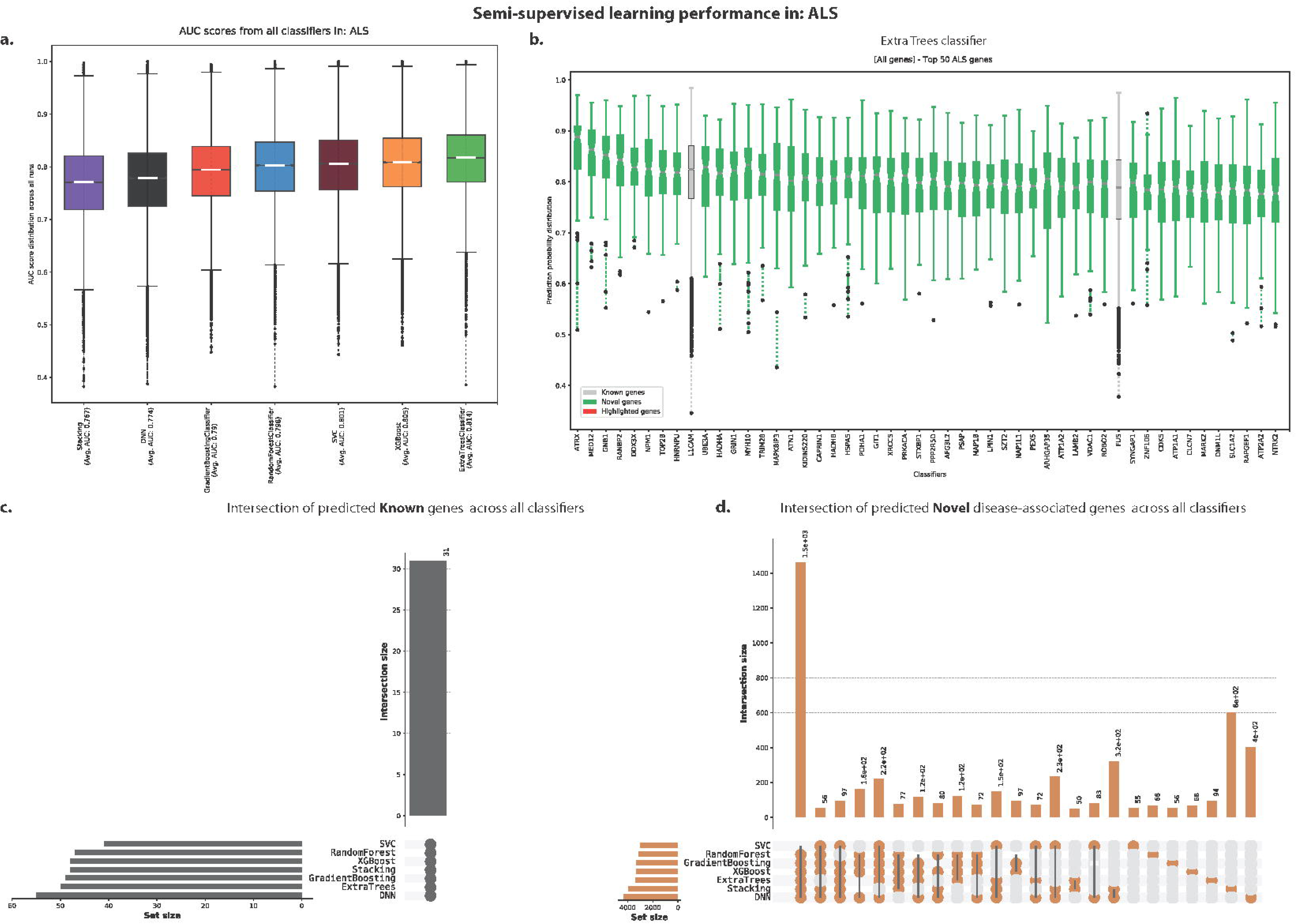
Mantis-ml performance on the ALS disease case, a) AUC score distribution per standard classifier used during mantis-ml training, b) Prediction probabilities from the top 50 (known and novel) genes predicted with Extra Trees as the standard classifier, c/d) Intersection sets of predicted known/novel genes across all classifiers.

**Suppl. Fig. 9.**
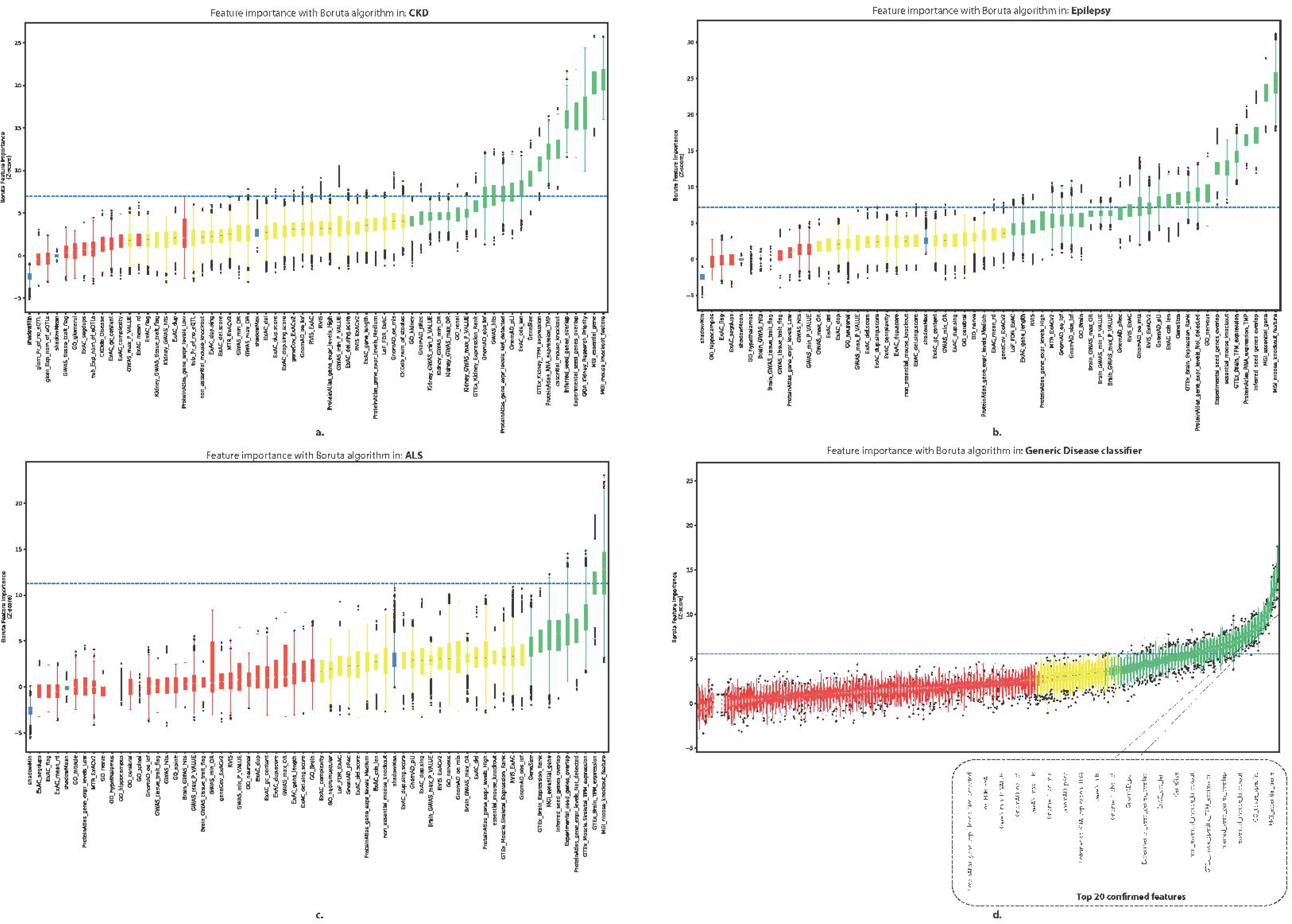
Distribution of feature importance scores extracted by a Random Forest classifier with the Boruta algorithm. Predictions are extracted across 100 balanced gene subsets with 10-fold cross-validation for the Chronic Kidney Disease (a), Epilepsy (b) and Amyotrophic Lateral Sclerosis (c) cases and for 1 balanced dataset with 10-fold cross validation for the Generic classifier (d). Confirmed features are shown in green, tentative in yellow and rejected ones with red. The random permuted features that are calculated as references by Boruta are shown in blue (’shadow’features). The top 20 confirmed features are shown for the Generic classifier.

**Suppl. Fig. 10.**
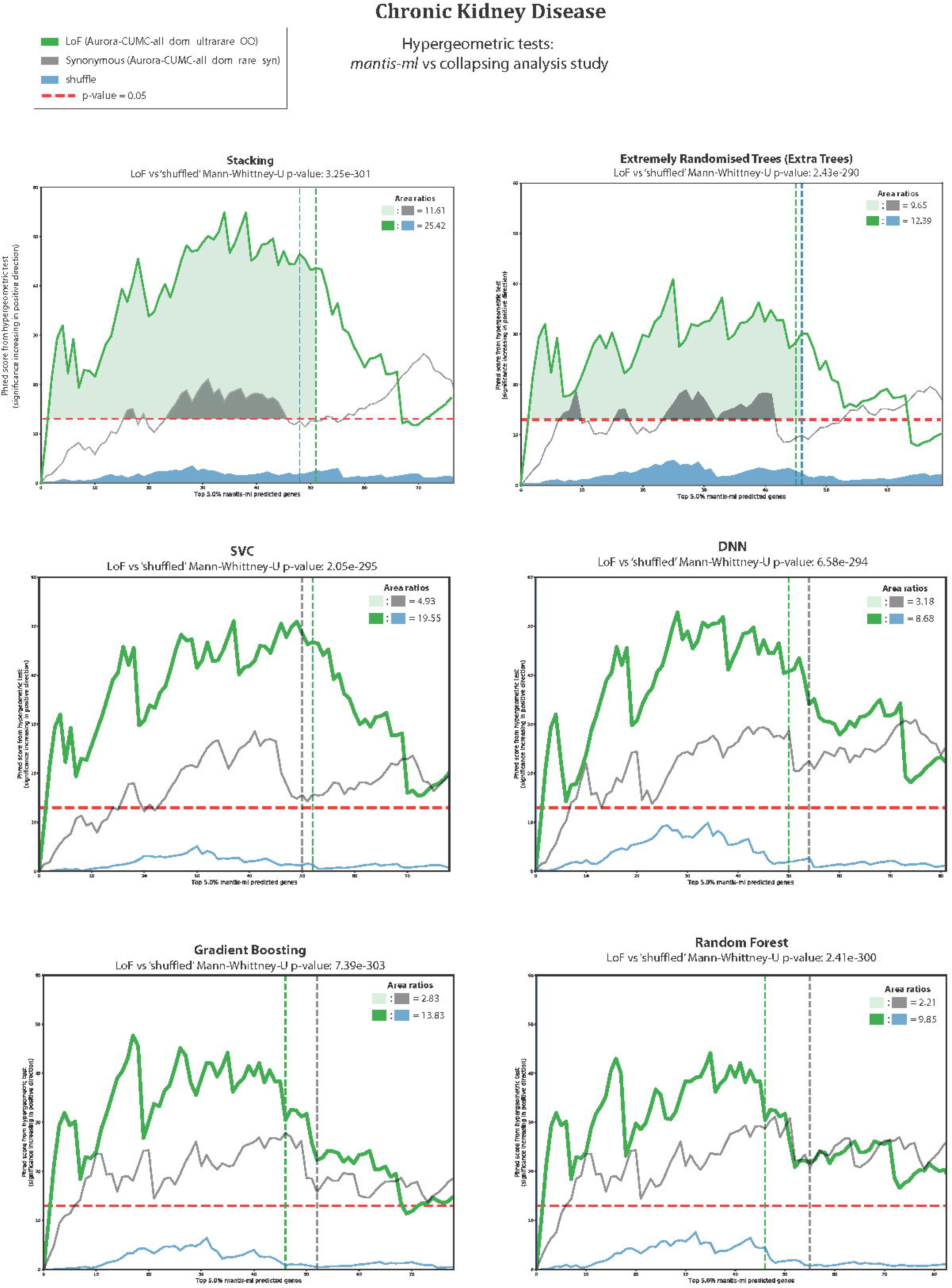
Cross-validation of mantis-ml predictions per classifier with rare-variant collapsing analysis results (applied for the Chronic Kidney Disease example)

**Suppl. Fig. 11.**
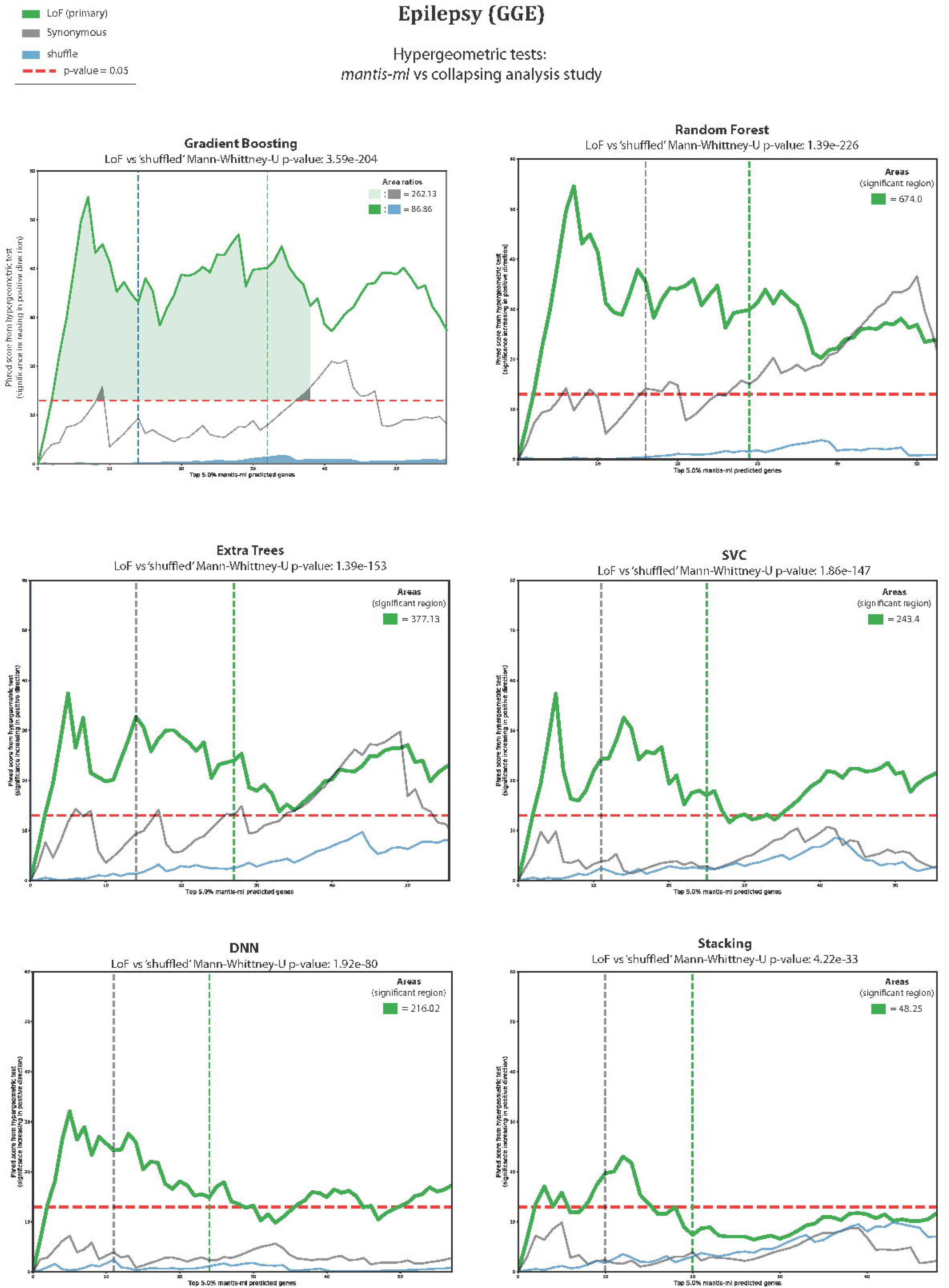
Cross-validation of mantis-ml predictions per classifier with rare-variant collapsing analysis results (applied for the Epilepsy disease example)

**Suppl. Fig. 12.**
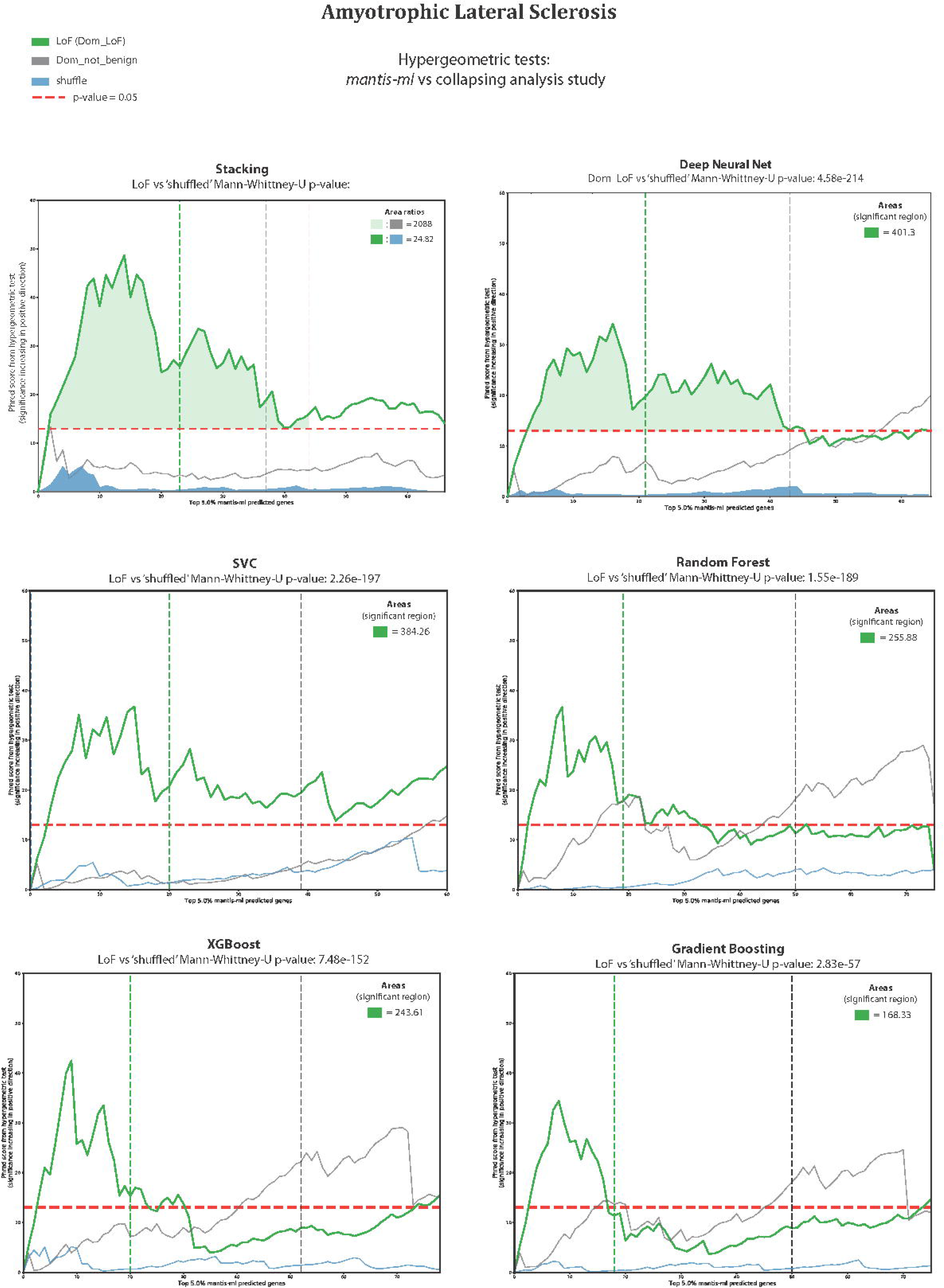
Cross-validation of mantis-ml predictions per classifier with rare-variant collapsing analysis results (applied for the ALS disease exampleh)

**Suppl. Fig. 13.**
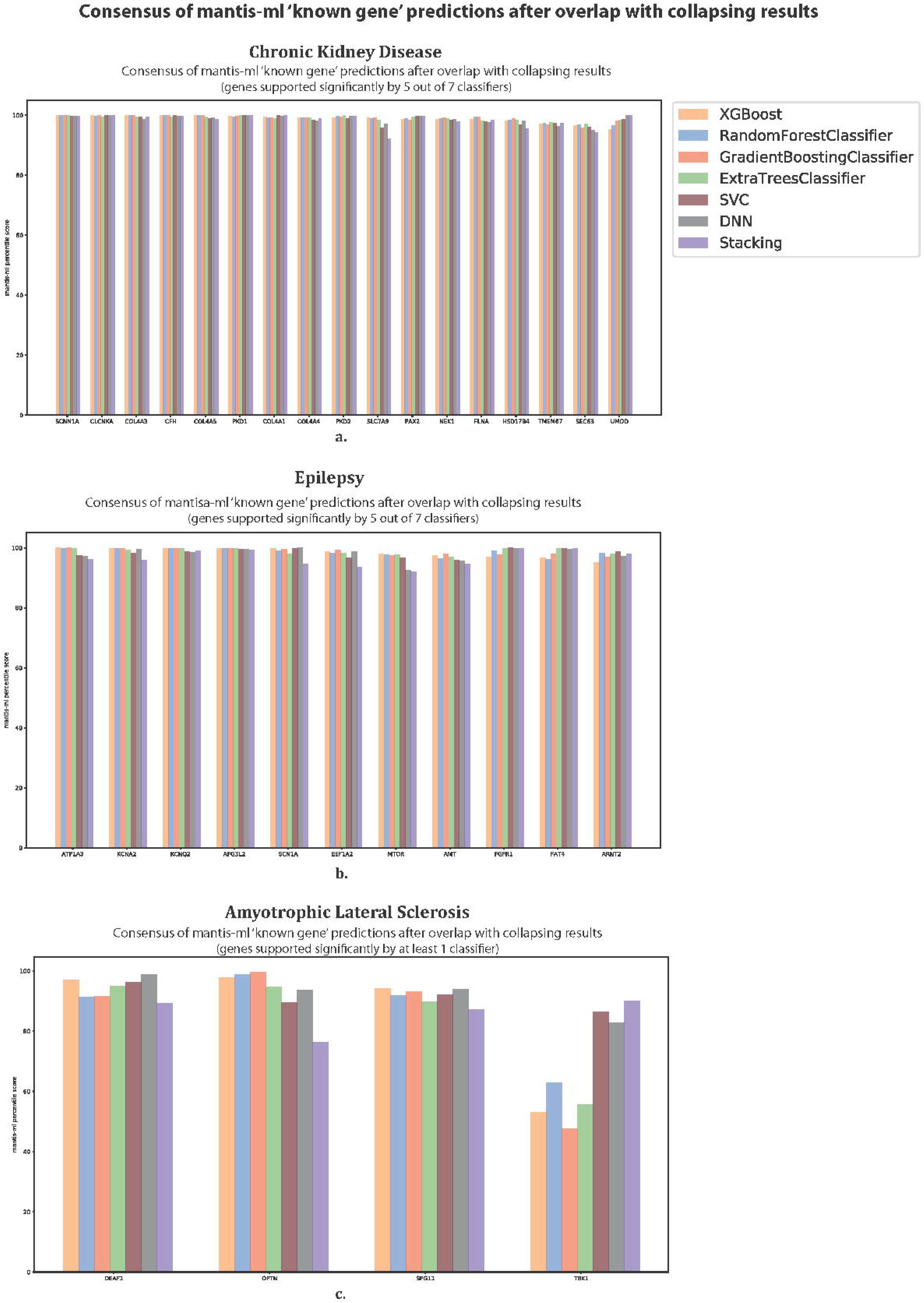
Consensus known genes across three disease cases (CKD, Epilepsy and ALS), satisfying the significance threshold criteria in both the collapsing analysis results and the hypergeometric, supported by multiple numbers of classifiers used by mantis-ml.

**Suppl. Fig. 14.**
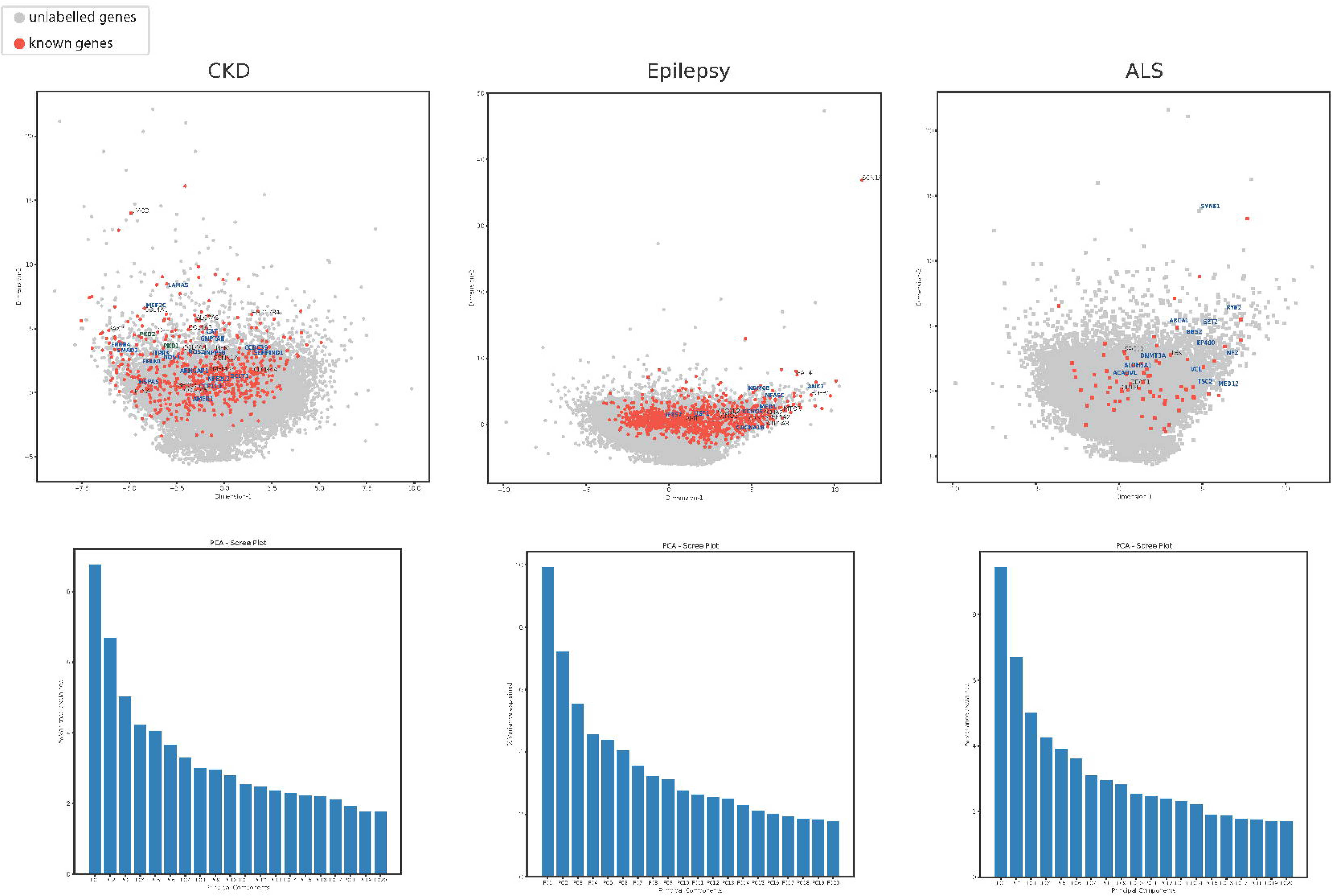
Principal Component Analysis (PCA) plots and Scree plots (variance explained by PCA components) for each disease example. Labelled genes are all the consensus novel (dark blue) and known genes predicted by overlapping mantis-ml predictions with rare-variant collapsing analysis results.

**Suppl. Fig. 15.**
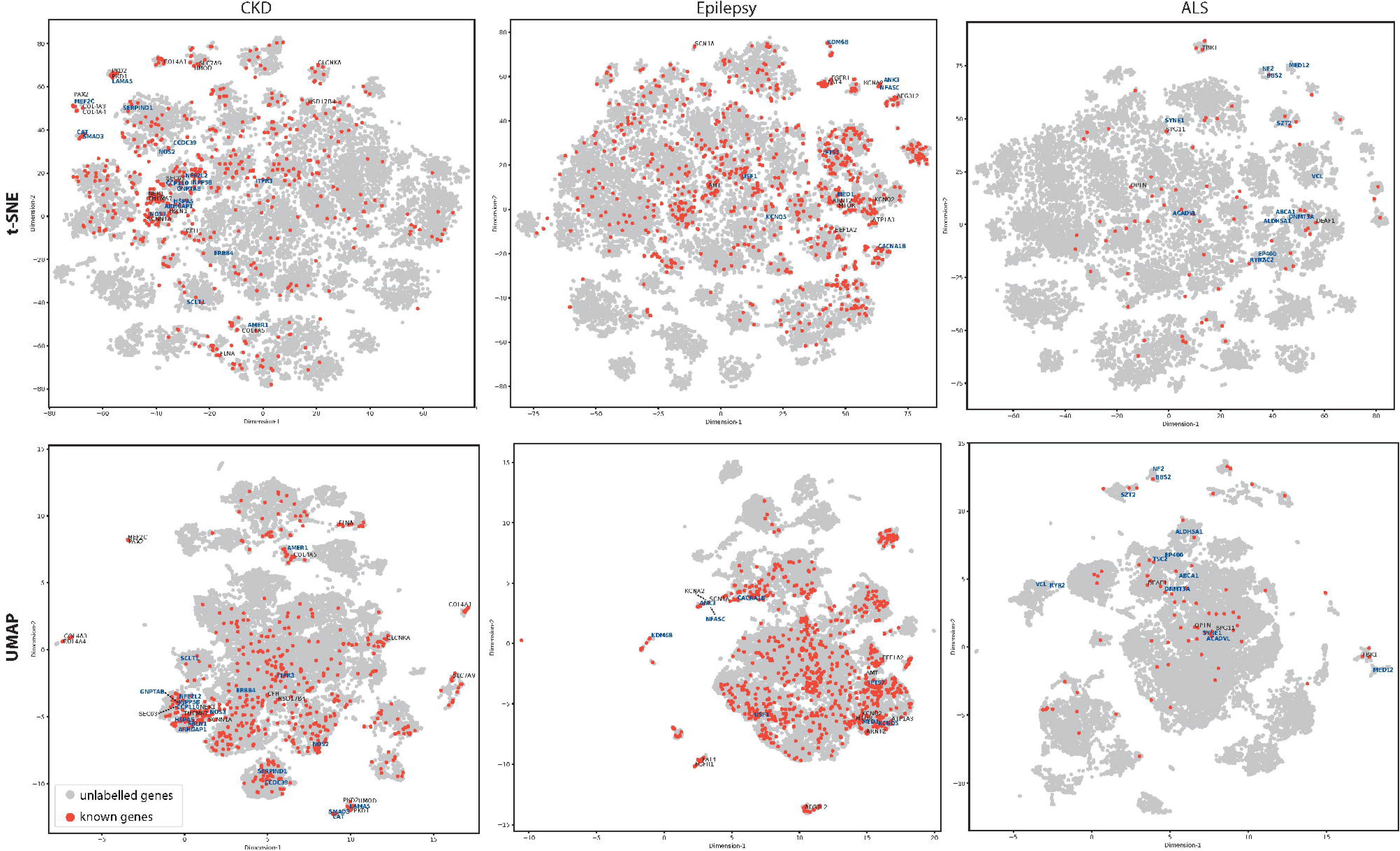
t-Distributed Stochastic Neighbor Embedding (t-SNE) and Uniform Manifold Approximation and Projection (UMAP) plots for 2D visualisation of all genes in each disease example. Labelled genes are all the consensus novel (dark blue) and known genes predicted by overlapping the mantis-ml predictions with rare-variant collapsing analysis results.

**Suppl. Fig. 16.**
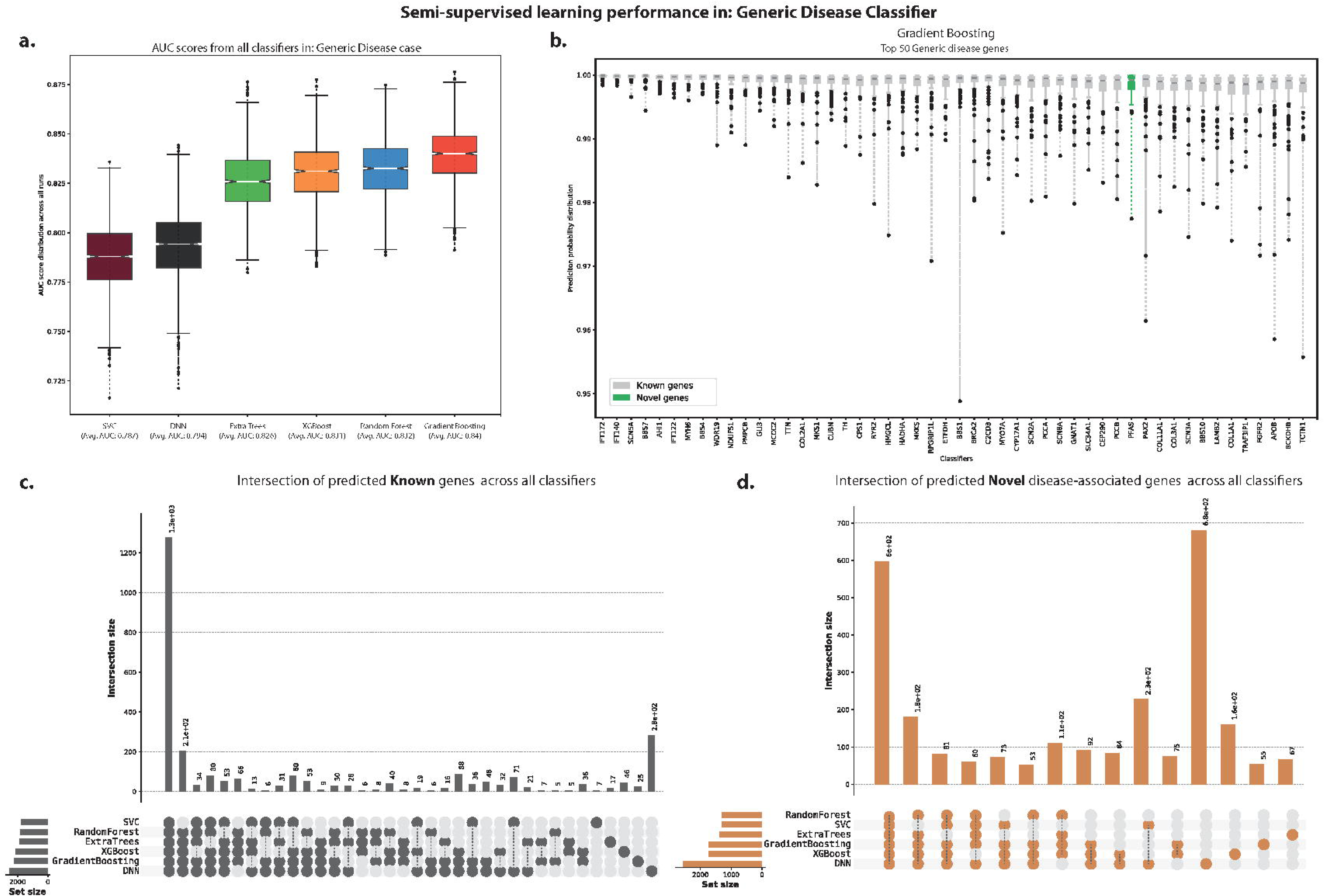
Mantis-ml performance on the Generic Disease case, a) AUC score distribution per standard classifier used during mantis-ml training, b) Prediction probabilities from the top 50 (known and novel) genes predicted with Gradient Boosting as the standard classifier, c/d) Intersection sets of predicted known/novel genes across all classifiers.

**Suppl. Fig. 17.**
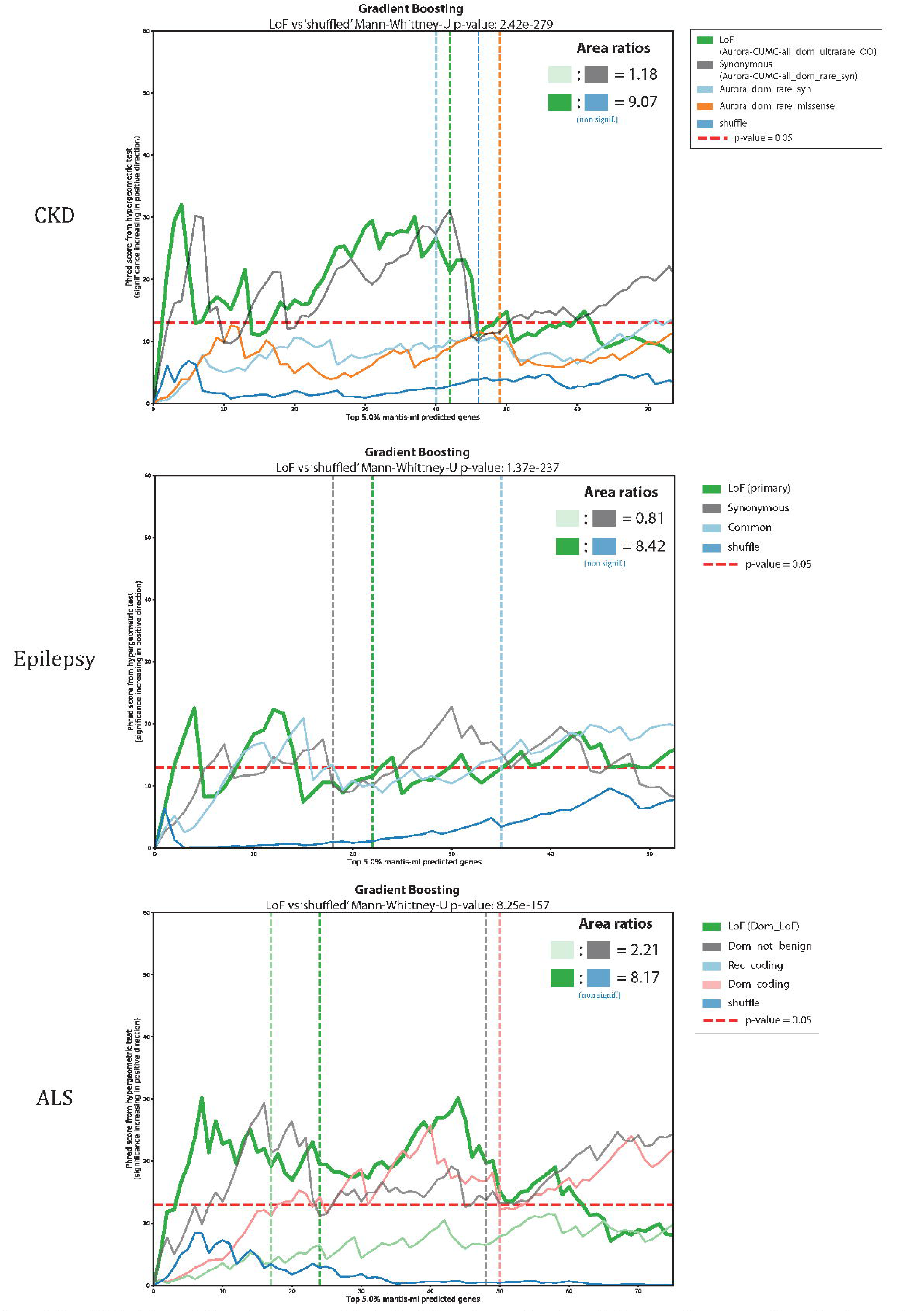
Enrichment (from hypergeo metric test) of Generic mantis-ml predictions on disease specific collapsing analysis results from CKD, Epilepsy and ALS related studies.

**Suppl. Fig. 18.**
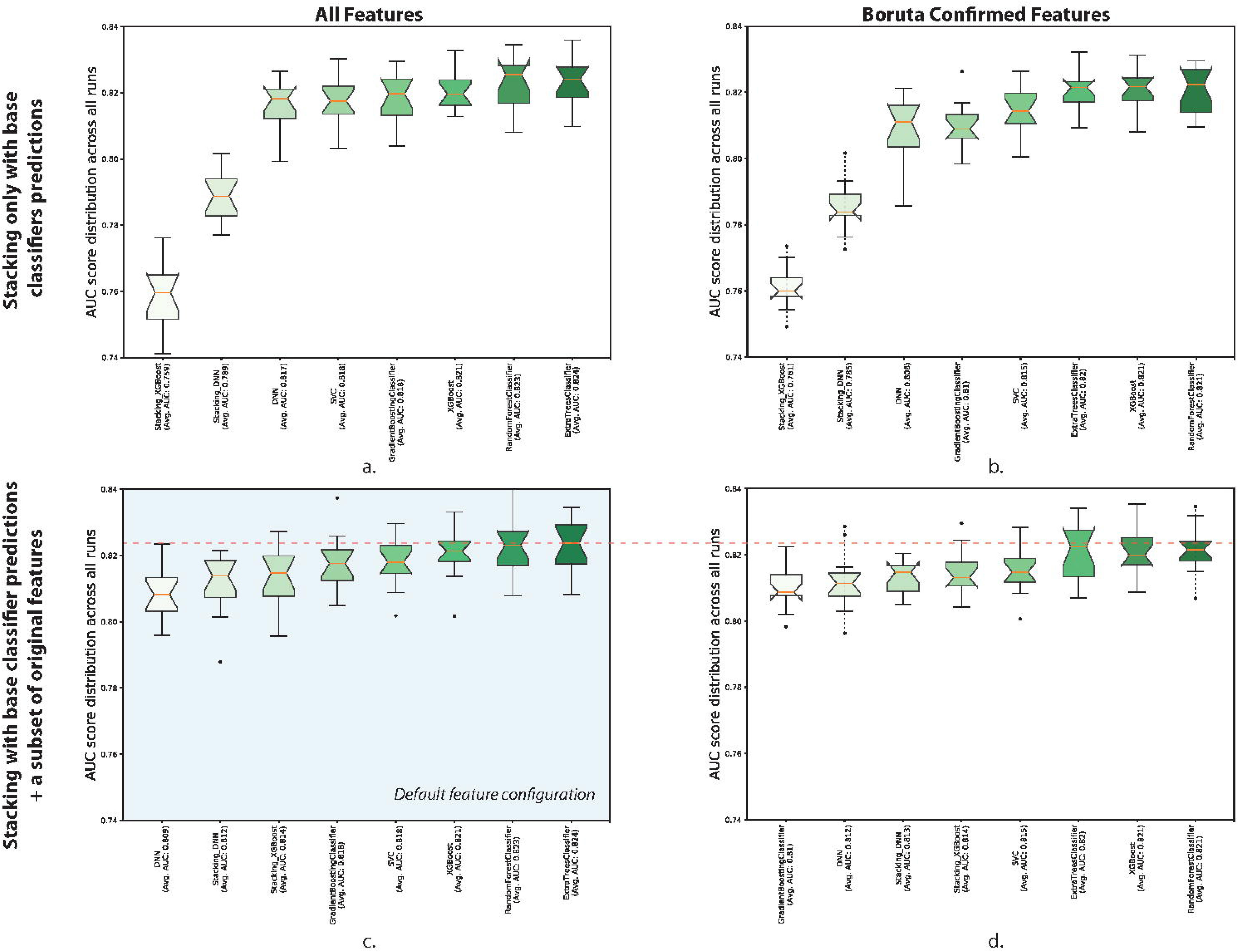
Benchmarking: AUC scores distribution from all classifiers in different feature selection configurations: a) using all features (after filtering) and training Stacking with base classifier predictions only, b) using only Boruta confirmed features and training Stacking with base classifier predictions + all Boruta confirmed features, c) using all features (after filtering) and training Stacking with base classifier predictions + all Boruta confirmed features and d) using only Boruta confirmed features and training Stacking with base classifier predictions + all Boruta confirmed features.Trainind data for benchmarking were compiled based on the Chronic Kidney Disease test case and included 15 random balanced datasets. Best average AUC performance was achieved by configuration (c) for all tested classifiers.

**Suppl. Fig. 19.**
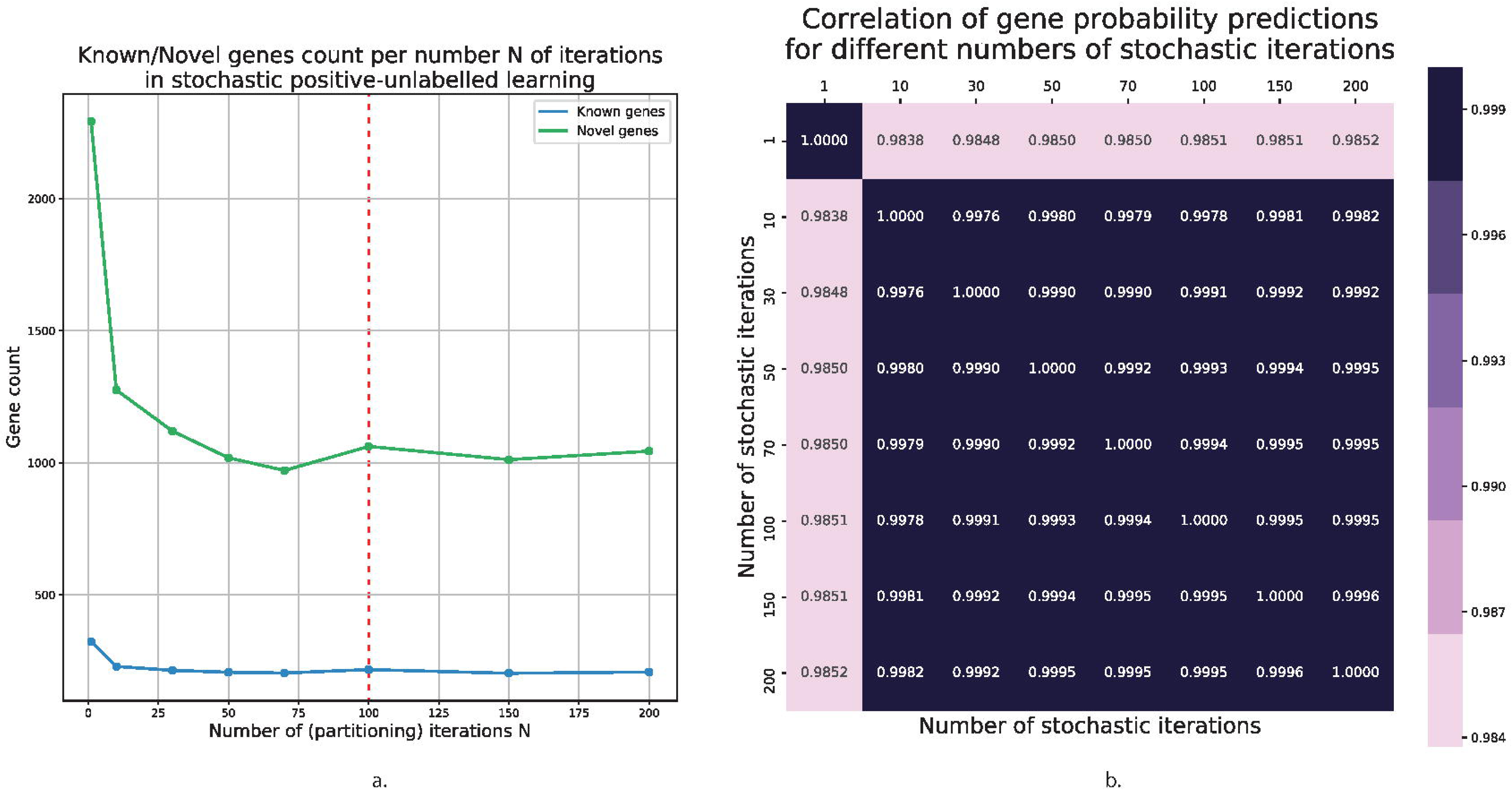
Benchmarking for: a) sensitivity check of the number of stochastic iterations to the predicted known and novel genes count and b) robustness of gene probability predictions for different numbers of iterations. (Tested numbers of iterations = [1,10,30,50,70,100,150,200]).

**Suppl. Fig. 20.**
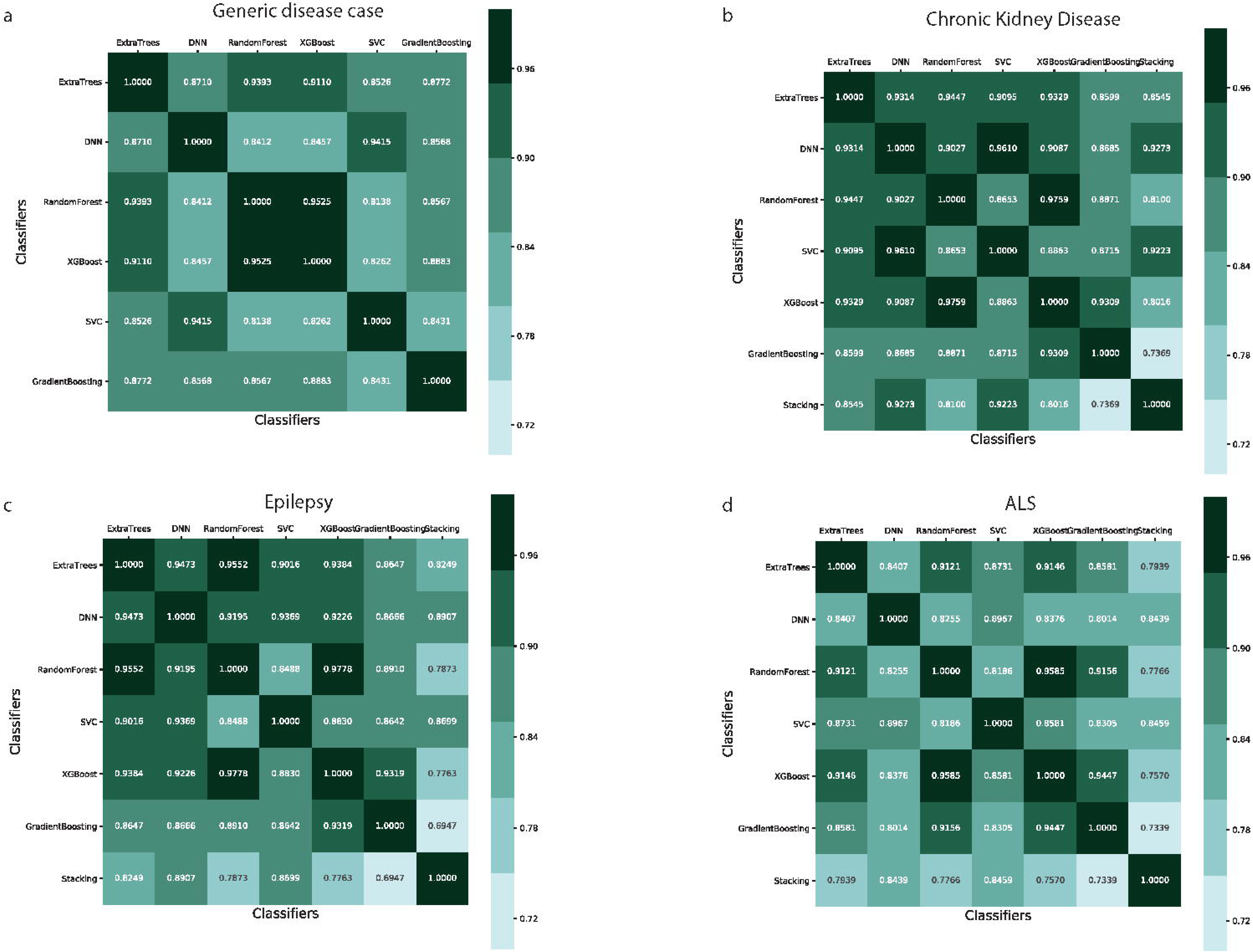
Correlation of gene probability predictions between different classifiers on the: a) Generic Disease, b) Chronic Kidney Disease, c) Epilepsy and d) ALS disease examples.

**Suppl. Fig. 21.**
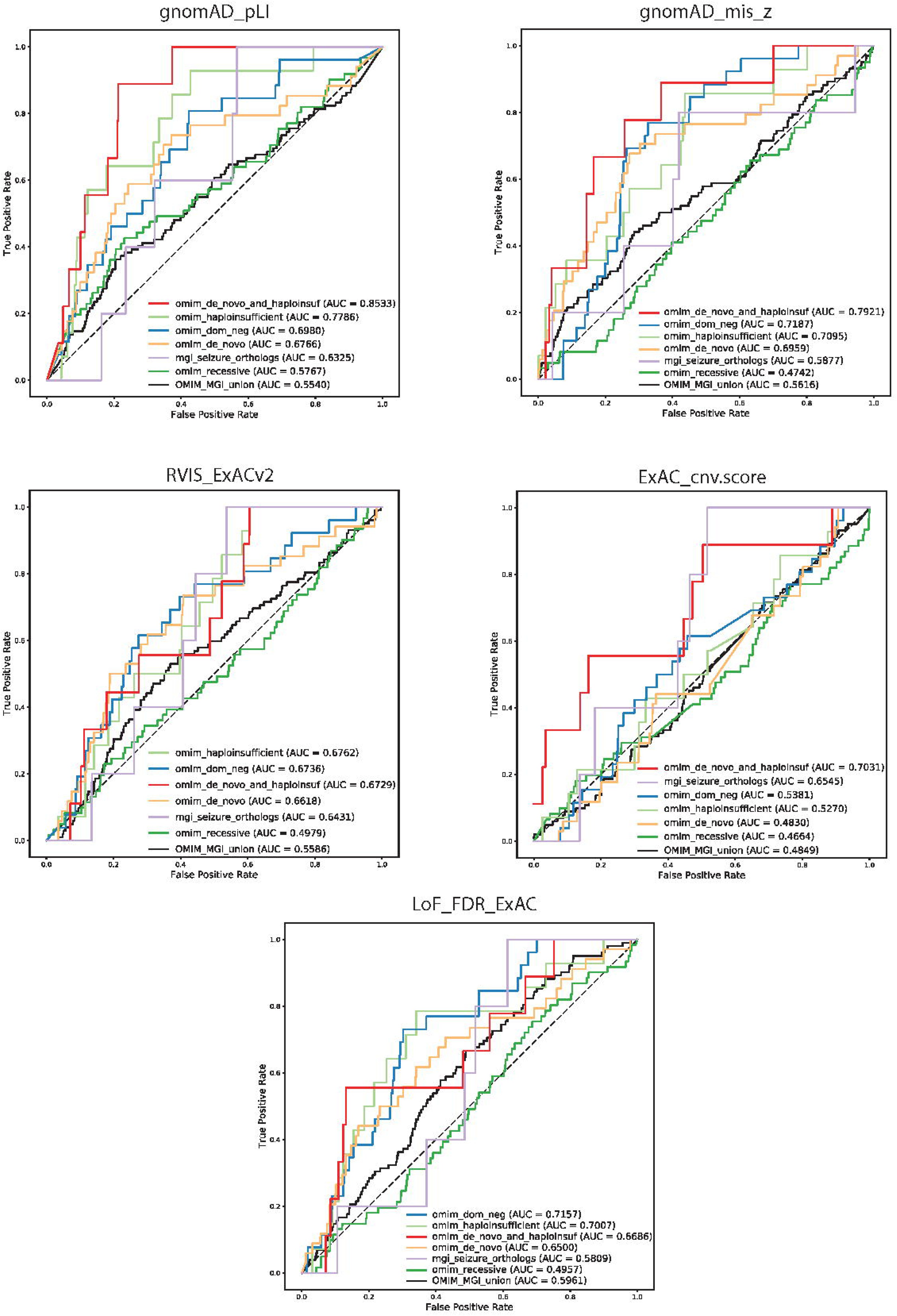
Predictive power of various intolerance scores for distinguishing different ΟΛΛΙΜ- and MGI-based genes from non-OMIM/MGI genes: gnomAD_pLI, gnomAD_mis_z, RVIS_ExACv2, ExAC_cnv.score and LoF_FDR_ExAC.

